# Monitoring oxidative inflammatory processes in live cells and tissue with Hypocrates, a genetically encoded biosensor for hypochlorite

**DOI:** 10.1101/2021.02.22.432222

**Authors:** Alexander I. Kostyuk, Maria-Armineh Tossounian, Anastasiya S. Panova, Marion Thauvin, Khadija Wahni, Inge Van Molle, Roman I. Raevskii, Mikhail S. Baranov, Sophie Vriz, Joris Messens, Dmitry S. Bilan, Vsevolod V. Belousov

## Abstract

Hypochlorous acid, an aggressive oxidant, is important in immune defense against pathogens. The current lack of tools to monitor the dynamics of hypochlorous acid in live cells and tissue hinders a better understanding of inflammatory processes. We engineered a genetically encoded biosensor, Hypocrates, for the visualization of hypochlorous acid. Hypocrates consists of a circularly permuted yellow fluorescent protein integrated into the structure of the transcription repressor NemR from *E. coli*. We determined sensitivity, selectivity, reaction rates, and the X-ray structure of this ratiometric redox biosensor, and tested the response of Hypocrates in HeLa Kyoto cells at varying hypochlorite concentrations. By combining Hypocrates with the biosensor HyperRed, we visualized the dynamics of hypochlorous acid and hydrogen peroxide in a zebrafish tail fin injury model.

## Introduction

In redox biology, much is known about the role of reactive oxygen species (ROS) in physiological and pathophysiological processes and about their cellular sources of generation. The best studied ROS is hydrogen peroxide (H_2_O_2_), which is not only cellular oxidative stress molecule but also a second messenger molecule that regulates cellular signal transduction pathways by modifying cysteine residues in proteins ^1–3^. Another oxidant, especially known to participate in immune response reactions is hypochlorous acid (HOCl). However, the role of HOCl remains one of the least explored areas. It is generated by the heme enzyme myeloperoxidase (MPO), as defense against bacterial infections ^4,5^. MPO catalyzes the conversion of Cl^−^ to OCl^−^ in the presence of H_2_O_2_. The formed HOCl can react with nucleophiles containing nitrogen and sulfur atoms, for example, with amines and thiols, as well as with aromatic rings in organic molecules. The possible cellular targets for HOCl are numerous: different amino acids in proteins, reduced glutathione, lipids, carbohydrates, and nucleobases ^6–8^.

Elevated activity of MPO and as a consequence increased levels of HOCl are often associated with diseases like atherosclerosis, diseases of the cardiovascular system and lungs, autoimmune diseases, Alzheimer’s disease, and many others ^9–12^. Although important, our knowledge on the spatial and temporal dynamics of HOCl is rather limited. It is still not known how HOCl participates in cellular signaling. For example, the exact role of N-chloramine, a milder and longer-lived oxidant compared to HOCl, that results from the reaction of amines with HOCl, is also not clear ^13–15^.

The most frequently used approaches to study hypochlorous stress in tissues is by measuring MPO enzymatic activity using colorimetric methods or immunohistochemical visualization of the enzyme localization. In addition, mass spectrometry and gas chromatography allowed the identification of chlorinated compounds as a result of exposure to HOCl. A drawback of these approaches is that they do not monitor real-time dynamics within live cells. Therefore, over the past few years, the market has seen a large number of fluorescent dyes for measuring HOCl ^16–22^. Despite all the advantages of these dyes, genetically encoded biosensors based on fluorescent proteins started revolutionizing redox biology research ^23^. Their main advantage over synthetic dyes is the ability to register the studied parameter in living systems of any level of complexity and this in real time. In particular, probes of the HyPer family contributed to our understanding of the biological role of H_2_O_2_^24–28^ Today, there is evidence that the aggressive oxidant HOCl also is involved in the regulation of proteins, modifying specific amino acid residues ^29–33^. However, the development of an indicator of protein nature for the visualization of HOCl seemed to be an impossible task.

Here we engineered a genetically encoded biosensor, the first of its kind for the visualization of HOCl in live cells and *in vivo* models. The biosensor is based on circularly permuted yellow fluorescent protein (cpYFP) integrated into the modified structure of the *E. coli* transcription repressor NemR. Wild-type NemR is sensitive to reactive chlorine species, including HOCl and chloramines ^32,34^, as well as to some electrophiles ^35^. It has been suggested that the oxidation with reactive chlorine species leads to the formation of a reversible sulfenamide bond between Cys106 and Lys175, which induces a local minor change in the protein conformation ^32,34^. Therefore, to develop a biosensor specific for HOCl, we used NemR with only a single cysteine, Cys106 (NemR^C106^). The newly developed biosensor was named Hypocrates (from **<u>Hypoc</u>**hlorite **<u>Rat</u>**iom**<u>e</u>**tric **<u>S</u>**ensor). The name is consonant with the name of the "Father of Medicine", Hippocrates. This greatest ancient physician was one of the first to depict signs of inflammation and to reflect on the nature of this process.

## Results

### Hypocrates (NemR-cpYFP biosensor) architecture and design

We decided to start this study by looking for prokaryotic transcription factors which sense hypochlorite anions (ClO^−^). HypR and NemR were selected ^32,33^. For our study, a NemR mutant with all Cys residues substituted for Ser, except for Cys106 (NemR^C106^) ^32^, was used to avoid undesirable sensitivity for reactive electrophilic species (RES) and to minimize other nonspecific redox reactions. We then measured the second-order rate constants of NemR^C106^ and HypR by monitoring the change of their respective intrinsic tyrosine and tryptophane fluorescence with increasing NaOCl concentrations (**Supplementary Fig. 1**). We found that NemR^C106^ (~1.1×10^5^ M^−1^s^−1^) reacts 160-times faster compared to HypR (~670 M^−1^s^−1^), and exposure to H_2_O_2_ had no effect (**Supplementary Fig. 2**).

Based on these observations, we selected NemR^C106^ as the molecular platform to design a biosensor for ClO^−^ detection. NemR^C106^ consists of a DNA-binding and a sensory domain. The sensory domain has a flexible loop with the critical Cys106 located at the C-terminus (**Fig. 1A)**. Based on the proposed protein functionality and structural flexibility^32^ (**Supplementary Fig. 3**), we hypothesized that after introducing cpYFP, the flexible loop could serve as a molecular switch capable of altering the optical properties of cpYFP (**Fig. 1B)**.

**Figure 1.**
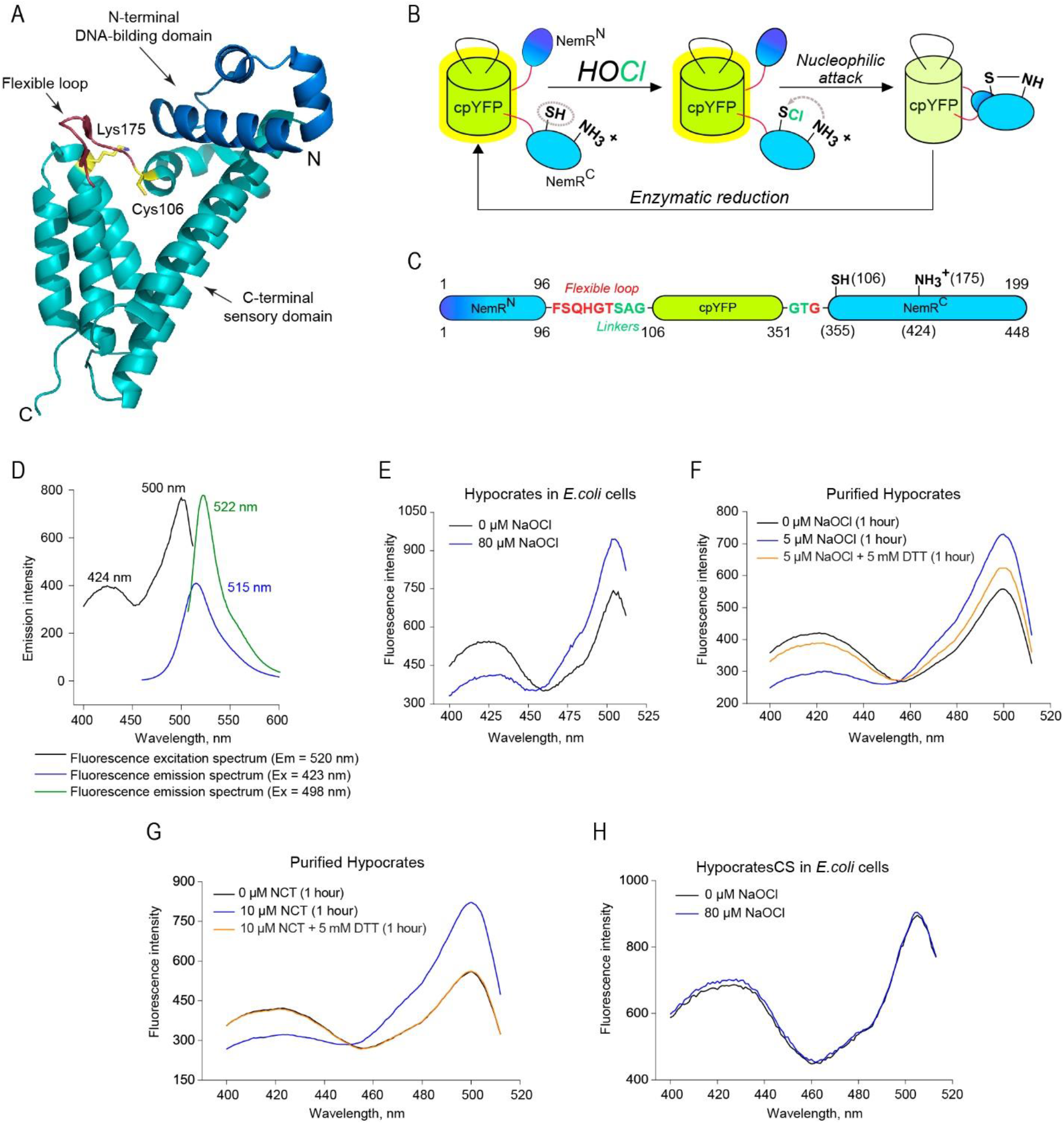
Hypocrates (NemR-cpYFP biosensor) design and spectral characteristics. (**A**) The structure of NemR^C106^ (PDB ID: 4YZE) shows the N-terminal DNA-binding domain (colored blue), the C-terminal sensory-domain (colored cyan), Cys106 and Lys175 (colored yellow) and the flexible loop (colored red), where the cpYFP was inserted. The N-and C-termini are indicated with N and C, respectively. (**B**) The proposed simplified scheme of NemR-cpYFP biosensors functioning in living cells. (**C**) The structure of Hypocrates is presented with NemR^C106^ colored blue/cyan, cpYFP colored yellow, the linkers between NemR^C106^ and cpYFP colored green and the flexible loop colored red. The upper numbers represent amino acid numbering corresponding to the intact NemR^C106^, while the lower numbers represent numbering corresponding to the biosensor. (**D**) The optical properties of purified Hypocrates protein in PBS. (**E**) Hypocrates fluorescence excitation spectra in *E. coli* cells in reduced and NaOCl-oxidized forms. (**F**) Purified Hypocrates (0.5 μM) fluorescence excitation spectrum behaviors in the presence of NaOCl in saturating concentration. (**G**) Purified Hypocrates (0.5 μM) fluorescence excitation spectrum behaviors in the presence of NCT in saturating concentration. (**H**) HypocratesCS fluorescence excitation spectra in *E. coli* cells in reduced and NaOCl-oxidized forms.

We constructed 12 chimers by introducing cpYFP in several positions of the NemR^C106^ flexible loop and by using a variation of short linkers (SAG/G or SAG/GT) (**Supplementary Fig. 4A**). We suggested that shortening of the cpYFP integration region can lead to a better signal transmission from the sensory to the reporter unit of the sensor. Therefore, we designed four chimers with one or two amino acid deletions in the flexible loop.

We recombinantly expressed all chimers in *E. coli*, tested the changes in the fluorescence excitation spectrum by adding NaOCl to the bacterial suspensions and to the purified chimers (**Supplementary Fig. 4B**). We selected the version with a maximum response amplitude of approximately 1.6 for further studies and named it Hypocrates (**Fig. 1C) (Supplementary Fig. 4C**). Purified Hypocrates protein is characterized by two excitation maxima (~425 nm and ~500 nm) and one fluorescence emission (~518 nm) maximum (**Fig. 1D**). The estimated Hypocrates brightness is in the ~4400-13900 range, depending on both the excitation wavelength and the redox state, which is approximately 7%-22% of the EYFP brightness (**Supplementary Tab. 1**). Further, after added NaOCl to *E. coli* cells expressing Hypocrates, we observed a ratiometric change in the fluorescence excitation spectrum (**Fig. 1E**). Thus, the biosensor signal can be calculated as a Ex_500_/Ex_425_ ratio. Purified Hypocrates behaved similarly and the ratiometric response can be reversed in the presence of a reducing agent (**Fig. 2F**). Next, we decided to test whether the biosensor is also sensitive to the HOCl-derivative N-chlorotaurine (NCT). NCT is one of the most common derivatives of reactive chlorine species because of the presence of relatively high taurine concentrations in neutrophils ^36^. Also for NCT, Hypocrates resulted in a fully reversible ratiometric response (**Fig. 1G**).

If Hypocrates works according to the above-proposed principles, then substitution of the key Cys355 residue for a nonreactive Ser should disrupt its sensing mechanism. We created this mutant version and named it HypocratesCS. As expected, *E. coli* cells expressing HypocratesCS did not respond anymore to the addition of NaOCl (**Fig. 1H**).

### The selectivity of Hypocrates

We showed that Hypocrates is highly sensitive to HOCl and NCT. It is noteworthy that high concentrations of NaOCl (~100 μM), but not NCT, led to pronounced fluorescence quenching due to apparent protein damage, which further indicates that NCT reacts with the sensor in a more specific way (**Supplementary Fig. 5**).

In addition, we investigated whether global structural changes of Hypocrates occurred in the presence of NaOCl using circular dichroism (CD). After adding NaOCl to the biosensor, we observed an increase of the molar ellipticity [θ] at 208 nm and 222 nm and a decrease at 194 nm (**Fig. 2A**). Upon H_2_O_2_ addition, no CD spectral changes were observed (**Supplementary Fig. 6**). To test whether the optical shift could be restored, we incubated the NaOCl-oxidized Hypocrates with the reducing agent DTT. After 5 min incubation with DTT, the spectrum of the oxidized biosensor showed a similar pattern as the one of reduced form (**Fig. 2B**), indicating the reversibility of the structural changes.

**Figure 2.**
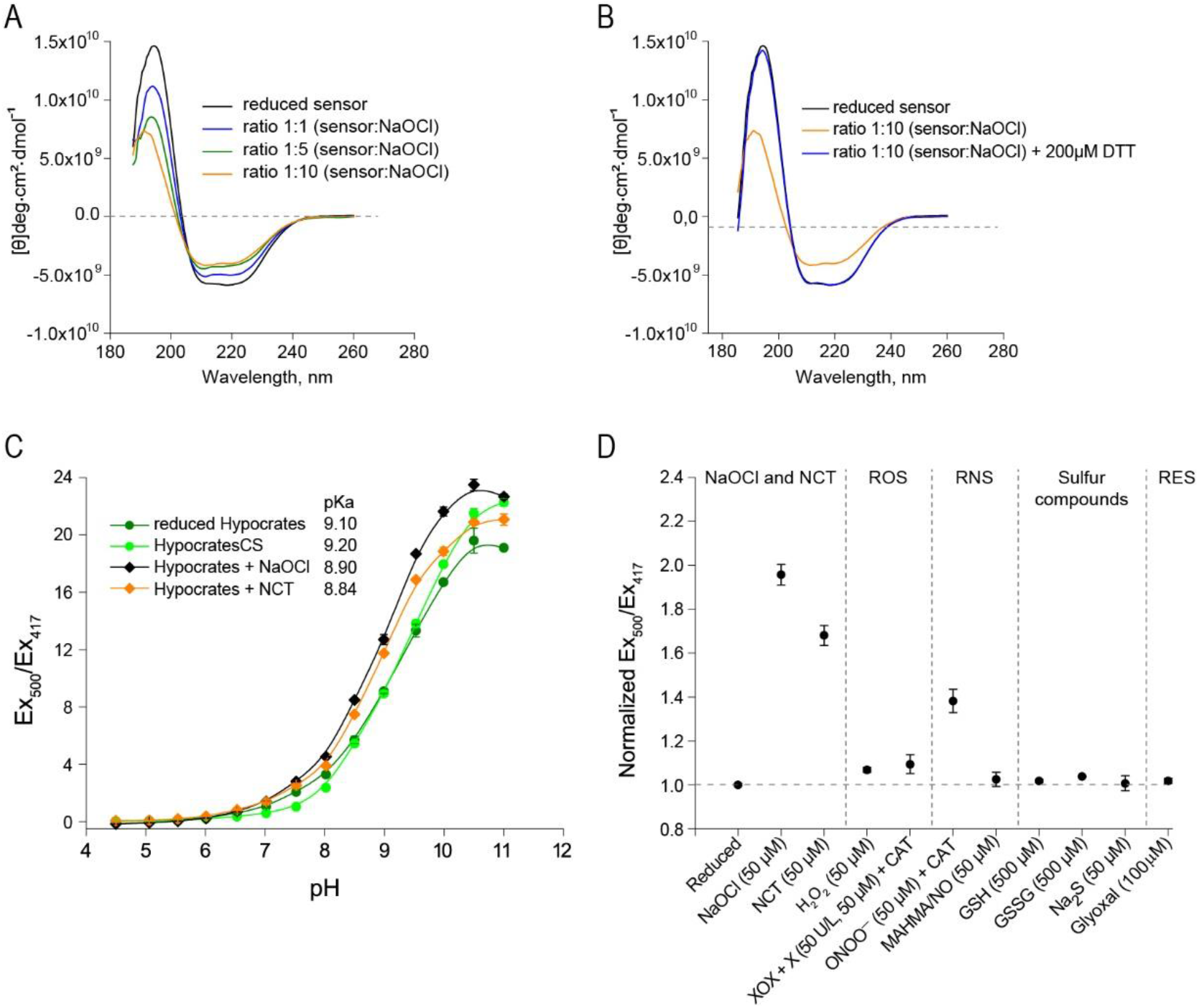
The selectivity of Hypocrates. (**A**) Far-UV circular dichroism spectra of reduced and NaOCl oxidized Hypocrates. With increasing NaOCl concentration, an increase of the molar ellipticity [θ] at 208 nm and 222 nm and a decrease at 194 nm were observed. (**B**) Upon reduction with DTT, the NaOCl-treated biosensor restores its overall secondary structure to the reduced form. (**C**) The excitation ratio of reduced and oxidized Hypocrates depends on the pH value of the buffer solution. The data are presented as a mean ± SEM, n ≥ 3. (**D**) Selectivity profile of purified Hypocrates towards a set of various redox compounds. ROS – reactive oxygen species, RNS – reactive nitrogen species, RES – reactive electrophilic species. Protein concentration was 2 μM in sodium/phosphate buffer in all samples except for the sample with glyoxal. In the sample with glyoxal, protein concentration was 0.5 μM in PBS. The data are presented as the mean ± SEM, n ≥ 3.

As for other cpYFP-based biosensors (except HyPer7 ^28^), the ratiometric response of Hypocrates is pH-dependent (**Fig. 2C**). The pKa of purified Hypocrates is 9.10, and HypocratesCS has a pKa of 9.20. In the presence of NaOCl and NCT, the pKa of Hypocrates decreases to 8.90 and 8.84, respectively. When changing the pH from 6 to 8, we observed a 12-fold signal increase; therefore, appropriate pH controls are required.

Next, we tested the selectivity of Hypocrates. We incubated the sensor with aliquots of various common oxidants (**Fig. 2D**). Only minor signal fluctuations were observed in the presence of high concentrations of H_2_O_2_, xanthine oxidase/xanthine system (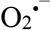 generator), MAHMA NONOate (NO^•^ generator), and GSSG (**Fig. 2D**). Although Hypocrates also shows a ratiometric response to ONOO^−^ (**Fig. 2D**) (**Supplementary Fig. 5**), the probability of such a response in biological systems is low because of the low sensitivity to ONOO^−^ and its low cellular concentration.

To validate whether the NaOCl-induced fluorescence changes are NemR^C106^-derived, we treated purified cpYFP (0.5 μM) and two other cpYFP-based biosensors (HyPer-2 ^25^ and SypHer3s ^37^) with NaOCl (5-10 μM) (**Supplementary Fig. 7**). cpYFP itself and also HyPer-2 and SypHer3s showed no response. As such, we concluded that cpYFP itself does not contribute to the ratiometric response. To obtain more direct evidence that the generated signal is NemR^C106^-derived, the intrinsic Trp fluorescence change of NemR^C106^ (2 μM) was determined in the presence of oxidizing agents (50 μM) (**Supplementary Fig. 7D**). Both NaOCl and NCT caused Trp-fluorescence changes (λ_ex_ = 295 nm, λ_em_ = 350 nm).

### Hypocrates sensitivity and reaction rates

The sensitivity of Hypocrates towards NaOCl and NCT was studied by titrating the biosensor (0.5 μM) with increasing oxidant concentrations up to 10 μM or 15 μM in sodium/phosphate buffer (**Fig. 3A-D**). In sodium/phosphate buffer, Hypocrates achieves saturation at ~4-5 μM (8-10:1 oxidant/sensor ratio). The Ex_500_/Ex_417_ ratio stabilizes at response values of ~1.8-fold (for NaOCl) and ~1.7-fold (for NCT) under saturating conditions. To estimate corresponding limits of detection (LOD), we implemented the 3S_y|x_/b approach, where S_y|x_ is the residual standard error and b is the slope of the linear regression model. In the described system, the LOD values are approximately 290 nM and 330 nM for NCT and NaOCl, respectively.

**Figure 3.**
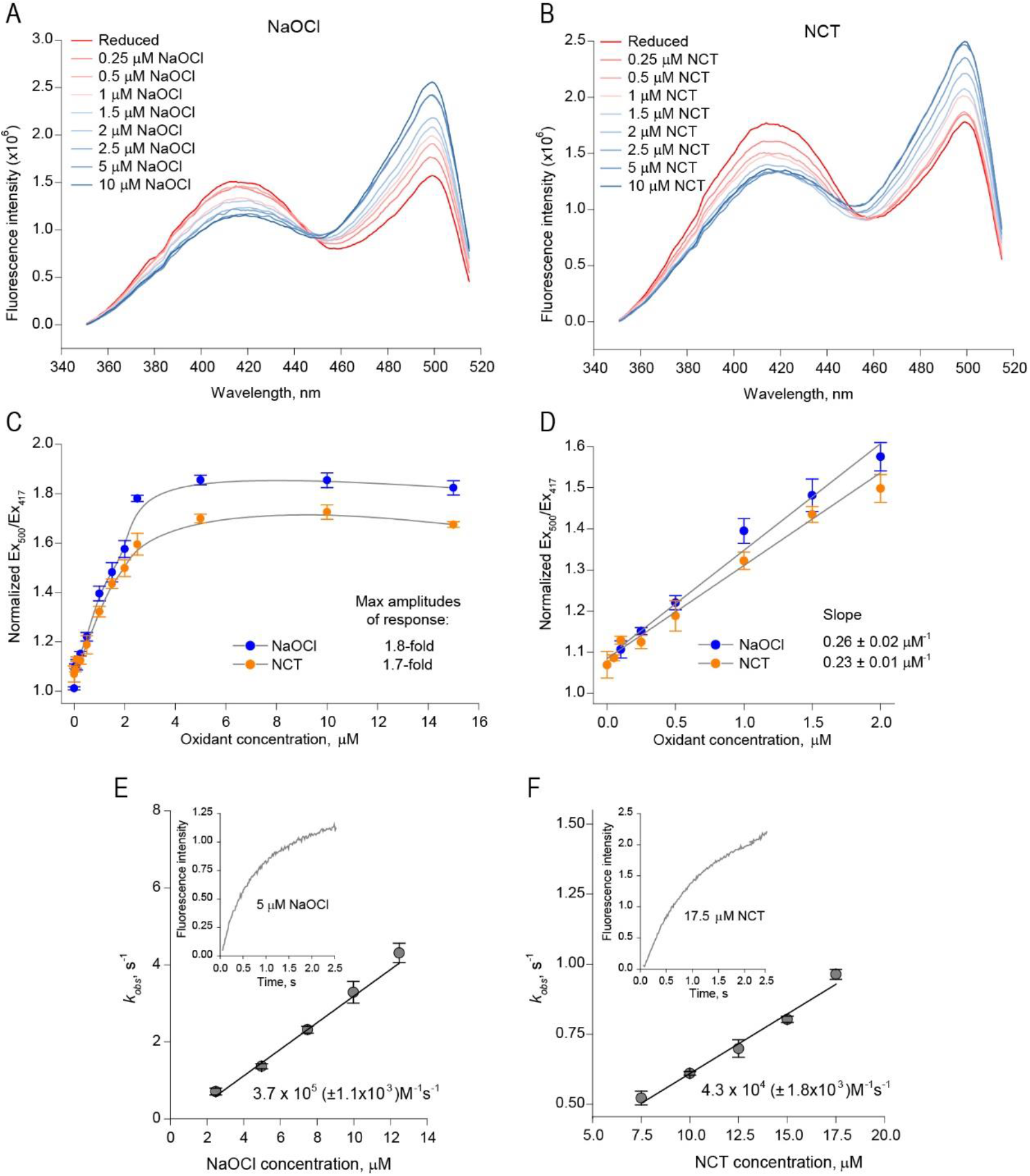
Hypocrates sensitivity and reaction rates. Changes in the fluorescence excitation spectra of Hypocrates (0.5 μM) obtained by sequential additions of (**A**) NaOCl or (**B**) NCT aliquots. (**C**) Titration curves of Hypocrates (0.5 μM) in sodium/phosphate buffer obtained by sequential additions of NaOCl or NCT aliquots. The data are presented as the mean ± SEM, n ≥ 2. The maximum amplitudes of response are 1.8-and 1.7-fold for NaOCl and NCT, respectively. In the presence of NaOCl and NCT, the probe is saturated at approximately 5 μM. (**D**) Hypocrates sensitivity towards NaOCl and NCT is shown. The data are presented as the mean ± SEM, n ≥ 2. (**E-F**) Hypocrates reaction rates. Changes in cpYFP fluorescence at >515 nm cut-off (λex = 485 nm) were measured as a function of time (insert). The curves were fitted to a single exponential to obtain the observed rate constants (k_obs_), which were plotted as a function of different (**E**) NaOCl or (**F**) NCT concentrations. The second-order rate constants of NaOCl (3.7 x 10^5^ (± 1.1 x 10^3^) M^−1^s^−1^) and NCT (3.4 x 10^4^ (± 1.8 x 10^3^) M^−1^s^−1^) were determined from the slope of the straight line. For each concentration, at least three independent experiments were performed.

To compare the reaction rates of Hypocrates towards NaOCl and NCT, the second-order rate constants were measured with the oxidants on a stopped-flow instrument (**Fig. 3E,F**). We found that the biosensor reacts faster with NaOCl (~3.7×10^5^ M^−1^s^−1^) compared to NCT (~4.3×10^4^ M^−1^s^−1^). NemR^C106^ is also less reactive to NCT compared to NaOCl (**Supplementary Fig. 8**), which possibly corresponds to the fact that NCT is a less aggressive compound.

### X-ray structure of HypocratesCS

To gain insights into the biosensor architecture and functional mechanism, we decided to crystallize Hypocrates and HypocratesCS. Only HypocratesCS gave diffraction-quality crystals. The orthorhombic crystals (C222_1_, a=90.242, b= 95.447, c=106.278, α=β=γ= 90°) contain one molecule of the biosensor per asymmetric unit and diffract to a resolution of 2.1 Å (**Supplementary Tab. 2**). The structure (PDB ID: 6ZUI) was solved by molecular replacement, using *E. coli* NemR^C106^ (PDB ID: 4YZE) and the cpYFP-based calcium sensor (PDB ID: 3O77) as search models. HypocratesCS consists of a NemR-based sensor domain (green) and a cpYFP domain (yellow) that undergoes a cyclization reaction to form the p-hydroxybenzylidene-imidazolidinone chromophore (orange), designated as “CR2” in the PDB (**Fig. 4A**). Superposition of the NemR^C106^ (PDB ID: 4YZE – blue) and HypocratesCS (sensor domain – green) shows a similar structure with a root mean square deviation (rmsd) of 0.506 Å for 159 atoms (**Fig. 4B).**As such, the insertion of cpYFP had only a minor structural effect on the overall structure of the NemR-sensor domain.

**Figure 4.**
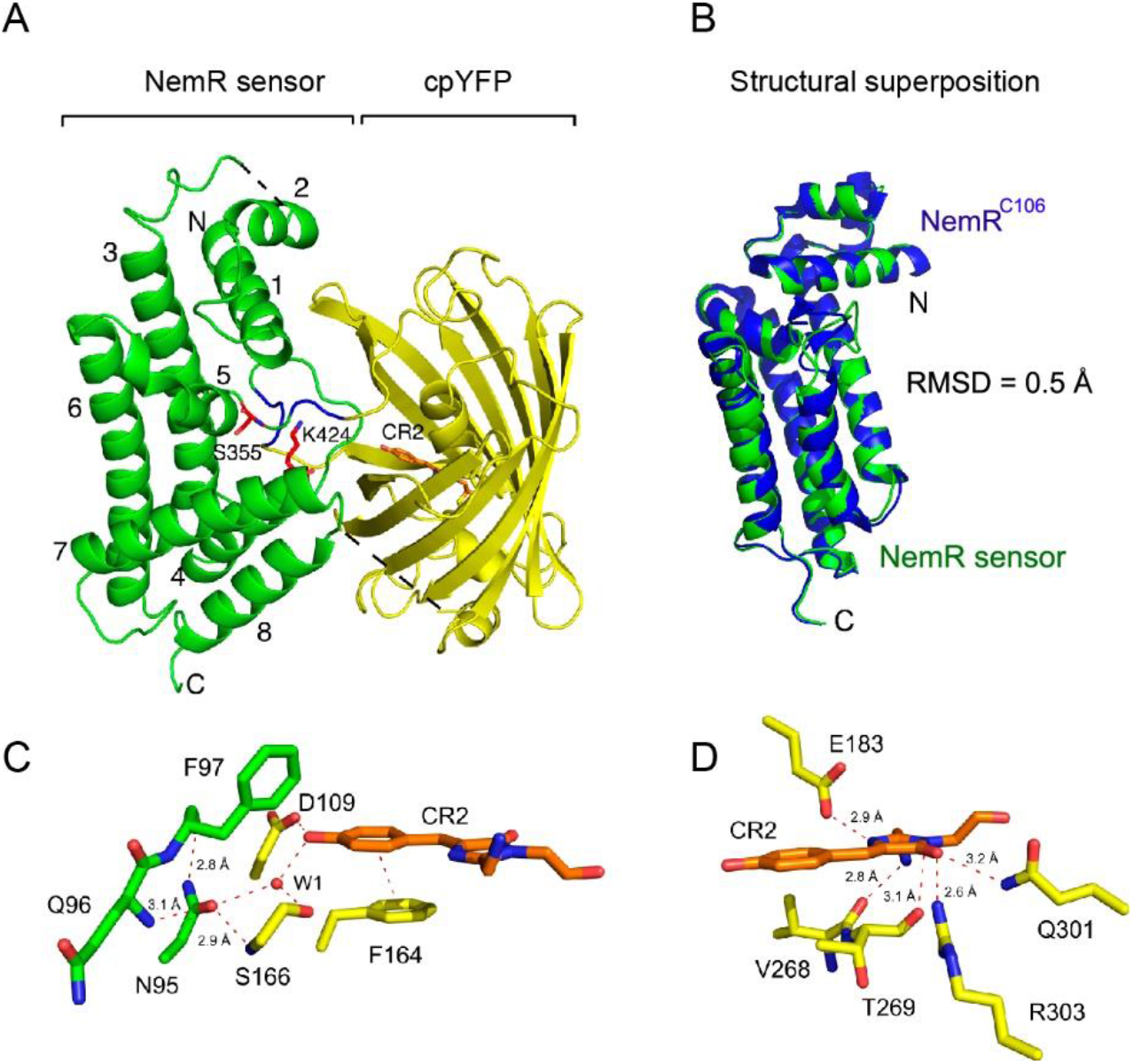
The structure of HypocratesCS, a cpYFP-based biosensor (PDB ID: 6ZUI). (**A**) The NemR-sensory domain (green) and the cpYFP domain (yellow) are shown. The chromophore (CR2) in the cpYFP β-barrel is shown in stick representation and colored orange. S355 and K424 are shown in stick representation and colored red. The linkers “SAG” and “GT” are colored blue. The missing segments (residues 40-42 and 191-207) are shown as black dotted lines. (**B**) Superposition of NemR^C106^ (blue – PDB ID: 4YZE) with the NemR-sensory domain (green). (**C**) N95 connects the sensory domain with cpYFP. N95 interacts with the backbone of Q96, F97 of the NemR-sensory domain and with S166 of the cpYFP domain. The 4-hydoxybenzyl group interacts the phenyl-ring of F164 over a distance of 3.9 Å. (**D**) The imidazolinone ring interacts with R303, Q301, V268, E183, and T269.

The structure of HypocratesCS is the first X-ray structure of a cpFP-based redox biosensor with an integrated cpYFP domain (**Fig. 4C**). The chromophore is in cis-configuration (**Supplementary Fig. 9**) and consists of an imidazolinone ring connected to a planar 4-hydroxybenzyl ring. The 4-hydroxybenzyl ring of the chromophore is stabilized by π-π stacking interaction with the phenyl-ring of Phe164, which has an off-centered parallel orientation to the 4-hydroxybenzyl ring. Aromatic ring stacking is also observed in YFP and in the cpYFP-based calcium sensor but is absent in GFP (**Supplementary Fig. 10C,D**) ^38–40^. The distance between the 4-hydroxybenzyl ring and the Phe164 phenyl ring is 3.9 Å. Further, the imidazolinone ring of the chromophore has several interactions with neighboring residues, suggesting that the chromophore will not change position during excited-state proton transfer (ESPT) (**Fig. 4D**).

Asn95 is the connecting residue between the NemR sensor domain and cpYFP (PDB ID: 6ZUI). Asn95 is located on the connecting loop between the α4 helix of the NemR-sensor domain and the N-terminus of the cpYFP domain. Asn95 interacts with the backbone of Gln96 and Phe97 in the NemR-sensor domain, and with Ser166, a conserved serine of the ESPT pathway in fluorescent proteins (**Supplementary Fig. 10A**). In the Ca^2+^ sensor, Case16, the domain connecting residue is Ser24 (**Supplementary Fig. 10C**) ^39^.

The ESPT pathways are different for cpYFPs, YFP, and GFP (**Supplementary Fig. 10A-D)**^38–40^ and consists of a hydrogen-bonding network surrounding the chromophore, a conserved serine and glutamate, and the conserved water molecules W1 and W2. In cpYFP (PDB ID: 6ZUI), the phenol oxygen of CR2 interacts with W1 and with Asp109, which might stabilize a negative charge on the phenol oxygen, similar to the phenol oxygen in GFP, where Thr203 takes over the role of Asp109 (**Supplementary Fig. 10A,D)**. Changing the position of Asn95 of the sensor domain could affect the pKa of the phenol oxygen via changes in the H-bond network in which W1, Ser166, and Asp109 are involved (**Supplementary Fig. 10A**), and this change could trigger a different charge transfer from the phenol oxygen via the phenol and imidazolinone rings with Glu183 as the final acceptor. The most likely final step of the pathway is a deprotonation of the heterocyclic ring nitrogen and the protonation of Glu183, rendering both groups neutral, but determining the exact details of the ESPT pathway and the potential role of W2 is beyond the scope of the present study.

### Hypocrates performance *in vitro* and in eukaryotic cell culture

We tested the ability of Hypocrates to visualize myeloperoxidase (MPO) activity *in vitro*. Incubation of the purified protein (0.5 μM) in the presence of the MPO-H_2_O_2_ system leads to a ratiometric response with an amplitude shift of 1.79-fold after 10 min of incubation, while H_2_O_2_ at a physiologically irrelevant high concentration (100 μM) induces only a minor oxidation shift of approximately 1.1-fold (**Fig. 5A,B**).

**Figure. 5.**
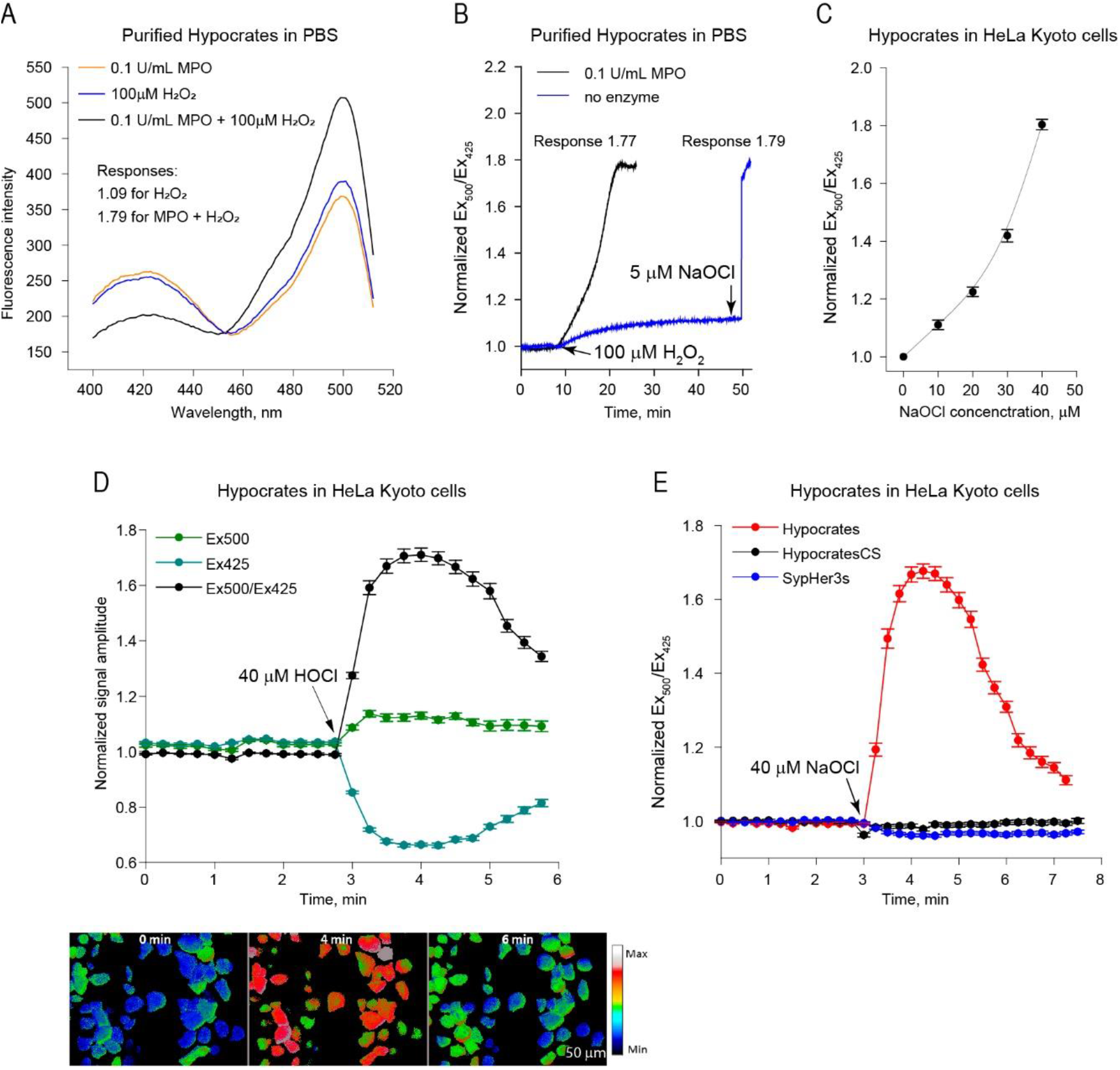
Hypocrates performance *in vitro* and in eukaryotic cell culture. (**A**) Fluorescence excitation spectra of purified Hypocrates (0.5 μM) in the presence of individual MPO, H_2_O_2_ and MPO-H_2_O_2_ system. (**B**) Hypocrates (0.5 μM) signal as a function of time in the presence of individual H_2_O_2_ and MPO-H_2_O_2_ system. HOCl, generated by MPO, leads to the development of a saturating response, while a physiologically irrelevant H_2_O_2_ concentration induces only minor signal changes. (**C**) The titration curve of Hypocrates in HeLa Kyoto cells exposed to different concentrations of NaOCl (values ± SEM, N = 2 experiments, n ≥ 25 cells per experiment). (**D**) Upper part: The timing of Hypocrates fluorescence changes induced by 40 μM NaOCl (values ± SEM, N = 2 experiments, n ≥ 30 cells per experiment). Lower part: Images of Hypocrates in transiently transfected HeLa Kyoto cells exposed to 40 μM of NaOCl at different time points. Scale bar = 50 μm. The lookup table indicates changes in the Ex_500_/Ex_425_ ratio. (**E**) The timing of Hypocrates, HypocratesCS and SypHer3s Ex_500_/Ex_425_ ratio changes after addition of 40 μM NaOCl (values ± SEM, N = 3 experiments, n ≥ 28 cells per experiment).

To test whether Hypocrates is functional in a eukaryotic system, we expressed the sensor in HeLa Kyoto cells and visualized the signal using fluorescence microscopy. To evaluate the sensitivity of the probe, we tested increasing concentrations of NaOCl and calculated the response as a Ex500/Ex425 ratio. The minimal concentration that induced detectable changes of the sensor fluorescence was approximately 10 μM NaOCl (**Fig. 5C**). Exposure to 40 μM NaOCl led to a signal change of 1.8-fold, which is similar to the saturating response obtained with purified protein and in *E. coli* suspension **(Fig. 1E,F)**. The oxidation of the biosensor in HeLa Kyoto cells was reversible – Hypocrates returned to the initial signal within approximately 3 min after NaOCl addition (**Fig. 5D**). We also transfected cells with HypocratesCS and with the specific pH-sensor SypHer3s ^37^. Exposure to 40 μM NaOCl did not significantly affect the signal of both probes (**Fig. 5E**), indicating that the response of Hypocrates, observed in this system, specifically reflects an NaOCl-induced response.

### Hypocrates performance in a zebrafish tail fin injury model

To test Hypocrates *in vivo*, we decided to induce inflammation using tail fin amputation of zebrafish larvae as a model. With the genetically encoded sensor HyPer ^24^, it was previously shown that the H_2_O_2_ concentration significantly increases in the wound margin and reaches its maximal value approximately 20 min post amputation ^41^. To obtain a full picture of the inflammation, we now decided to simultaneously monitor the H_2_O_2_ and OCl^−^ production. Therefore, we combined Hypocrates with HyPerRed ^27^, a red sensor for H_2_O_2_ (**Fig. 6**). Signals of both Hypocrates and HyPerRed increased 15 min post amputation (mpa), then HyPerRed fluorescence decreased while Hypocrates signal changed much more slowly. In parallel, we used the control version HypocratesCS. Although HypocratesCS signal also increased, statistical analysis revealed that the difference between the response of Hypocrates and HypocratesCS was significant. We demonstrated that Hypocrates is suitable for *in vivo* imaging with HypocratesCS as a control.

**Figure. 6.**
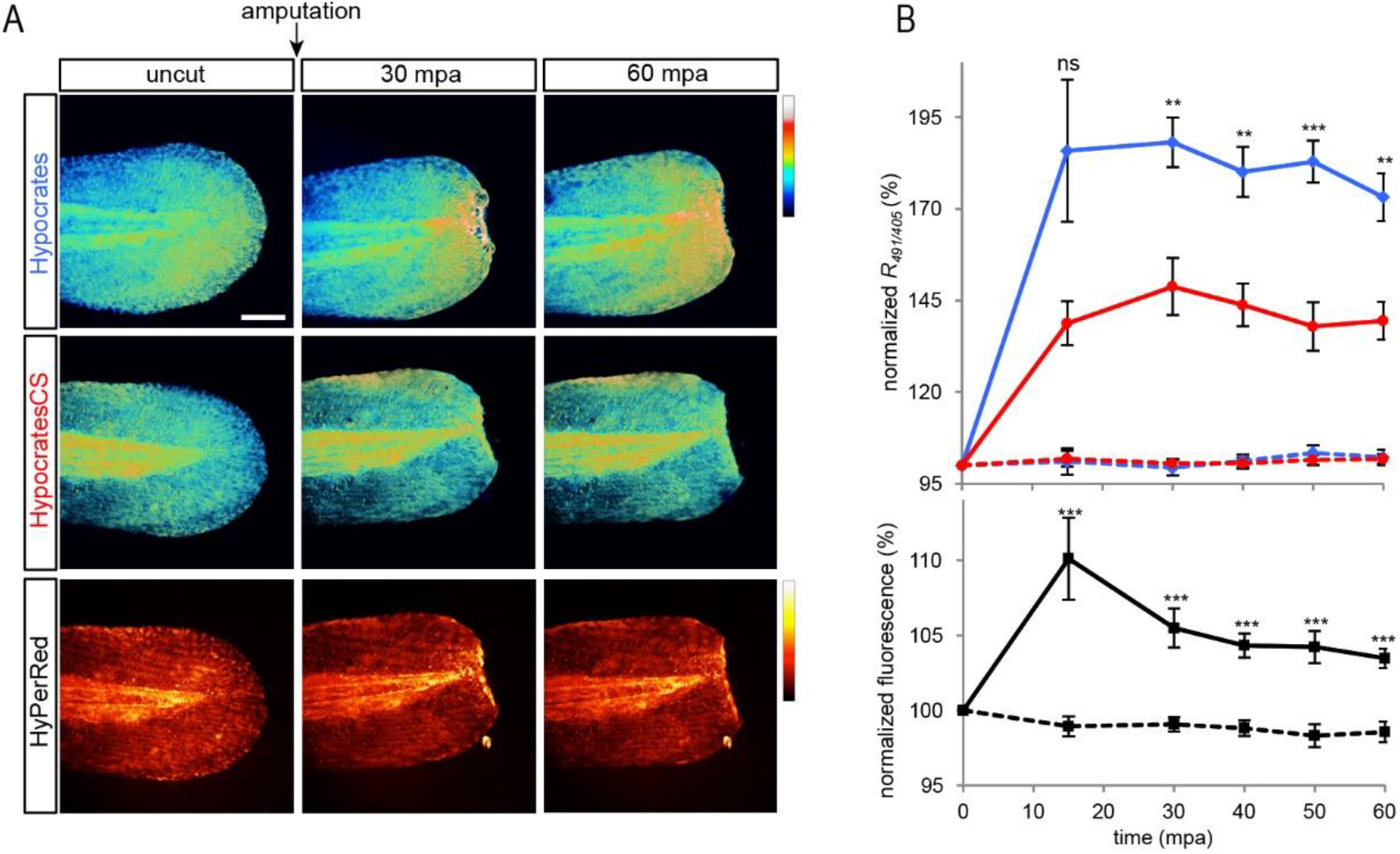
Hypochlorite and H_2_O_2_ dynamics during zebrafish larvae wounding. (**A**) Hypocrates and HyPerRed imaging. Zebrafish embryos were co-injected with Hypocrates or HypocratesCS and HyPerRed mRNAs at the 1-cell stage, and a tail fin amputation assay was performed on 48 hpf larvae. Images were taken before amputation, and time lapse imaging was performed up to 60 min post amputation (mpa). Scale bar = 100 μm. (**B**) Hypocrates ratio and HyPerRed fluorescence were quantified at the amputation plane and normalized to the mean fluorescence on uncut tail for each larva. Ratio quantification on larvae tail fin expressing Hypocrates (blue lines) or HypocratesCS (red lines). Non-amputated embryos (dashed lines) expressing Hypocrates or HypocratesCS were also imaged as a control (values ± SEM; N = 4 experiments, n ≥ 3 embryos/timepoint; ns, no significant, **, *P* < 0.01, ***, *P* < 0,001, *versus* HypocratesCS cut larvae). HyPerRed fluorescence quantification on larvae tail fin expressing HyPerRed (black lines) (values ± SEM, N = 3 experiments, n ≥ 7 embryos/timepoint; ***, *P* < 0,001, *versus* uncut larvae (dashed line)).

## Discussion

HOCl imaging can be addressed with a quite extensive set of low molecular weight dyes with marked variability in optical properties and chemical structure of the sensing moiety ^16–22,42,43^. Although these dyes often provide sufficient selectivity and were successfully implemented in cell cultures as well as in whole organisms, they are characterized by a large number of technical disadvantages. The quantitative interpretation of the data obtained with these dyes is significantly hampered since the vast majority of these chemicals generate an intensiometric response. Long term visualization of repetitive redox events also seems to be problematic due to the irreversible nature of HOCl-induced modifications that underlie the sensing mechanisms of most of these dyes. Further, the reaction rates of the current HOCl dyes are relatively low and signal stabilization usually requires dozens of seconds or even minutes ^44^; in particular, the measured rate constants for CMOS and FDOCl-1 are ~0.67 s^−1^ and ~0.10 s^−1^, respectively ^45,46^. As a consequence, rapid shifts in HOCl concentrations will not be visualized, and most likely will lower the actual sensitivity because of the presence of kinetically more favorable HOCl targets of the cell. Finally, compared to genetically encoded tools, fluorescent dyes are characterized with a low spatial resolution. Based on the drawbacks of current tools and on the fact that more and more proteins are characterized as being modified under hypochlorous stress conditions, the idea of creating a genetically encoded biosensor for visualizing hypochlorous acid and its derivatives was born.

We present the Hypocrates probe, which is the first genetically encoded fluorescent biosensor for visualizing HOCl in live systems. Hypocrates displays a ratiometric reversible change in signal when interacting with HOCl and NCT with minimal response-inducing oxidant concentrations in the 0.1-0.3 μM range (at the biosensor concentration of 0.5 μM). It is known that neutrophils produce high amounts of HOCl. It is difficult to calculate the exact concentration of HOCl produced by cells since it quickly reacts with surrounding molecules. However, the concentration of HOCl in the interstitial fluids of inflamed tissues has been estimated to reach several millimolar ^47^. Hypocrates also allows monitoring the dynamics of HOCl derivatives. Chloramines are characterized by longer lifetimes due to decreased reaction rates and altered selectivity profiles with higher specificity for sulfur-containing groups. Due to the high concentrations of taurine in neutrophils, N-chlorotaurine is one of the most common derivatives of reactive chlorine species ^36^.

Recombinantly expressed and purified Hypocrates did not show any response to the major common intracellular oxidizing agents. However, we observed spectral changes of Hypocrates in the presence of ONOO^−^. The saturating concentration of ONOO^−^ for the maximum response of Hypocrates *in vitro* was 10 μM at a biosensor concentration of 0.5 μM. However, ONOO^−^ formation in cells directly depends on NO^•^, the concentration of which during physiological processes *in vivo* can range from 100 pM (or below) up to ∼5 nM ^48^. Although, in some cases it has been shown that NO^•^ reaches micromolar levels in tissues during specific pathological processes ^49^. The rate constant for the reaction of 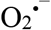 with NO^•^ to yield ONOO^−^ is one order of magnitude higher than 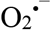 dismutation catalyzed by superoxide dismutases ^50^. Obviously, maximal generation of ONOO^−^ and its steady-state concentration should be achieved at sites with maximal production of 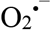 with NO^•^. However, equimolar fluxes of precursors, suggesting maximal formation of ONOO^−^, is a simplification of real events in biological systems. For this reason, the actual achievable concentrations of ONOO^−^ are difficult to calculate ^51^. In addition, in conditions of inflammation, many factors are in play, their relationship remains to be studied. For example, MPO in neutrophils may inhibit NO^•^ production via the formation of chlorinated L-arginine, which inhibits all types of NO^•^ synthases ^52^. However, for more accurate measurements various inhibitors of NO^•^ synthases can be used in control series of experiments using Hypocrates as biosensor.

The crystal structure of HypocratesCS is the first cpFP-based redox biosensor that reveals its CR2 chromophore environment within its overall structure (**Fig. 4A,C**). Overall structure comparison of the β-barrel of cpYFP in Hypocrates (PDB ID: 6ZUI) with the calcium biosensor, Case16 (PDB ID:3O77) ^39^, showed that both β-barrels are very similar (rmsd of 0.282 Å for 190 atoms). Further, both have a CR2 chromophore, while GFP is characterized by a CRO chromophore (**Supplementary Fig. 10D)**. As such, the cpFP in Case16 is actually a cpYFP, and not a cpGFP as mentioned in Leder *et al*. ^39^. The structure of HypocratesCS is important not only for understanding the functioning of this biosensor but also for revealing the features of other cpYFP-based probes, which show subtle differences in their CR2 chromophore environment (**Supplementary Fig. 10A,C**).

All in all, Hypocrates is suitable for the study of inflammatory reactions *in vivo*. Here, we induced inflammation by injuring the caudal fin of *Danio rerio* larvae. Previously, it was shown with the HyPer biosensor that an H_2_O_2_ gradient is formed in the wound, which serves to attract neutrophils to the area of inflammation ^41^. Moreover, neutrophils subsequently participate in the elimination of the H_2_O_2_ gradient due to the reaction catalyzed by MPO ^53^. Here, we observed for the first time in multiparameter microscopy mode the simultaneous real-time dynamics of H_2_O_2_ and HOCl *in vivo* in zebrafish tissues during inflammation using the red fluorescent biosensor HyPerRed ^27^ and the green emitting Hypocrates.

## Data availability

The X-ray crystal structure of HypocratesCS was deposited in the protein data bank under accession code 6ZUI.

## Acknowledgements

The work was supported by the Russian Foundation for Basic Research (RFBR) Grant 18-34-20032 (to D.S.B.); the Russian Science Foundation (RSF) Grant 17-15-01175 (to D.S.B) for work related to the preparation and testing of Hypocrates biosensor in the eukaryotic system; the Grants from the Vlaams Instituut voor Biotechnologie (to J.M.); a FWO Ph.D. fellowship grant (to M-A.T.); CNRS, INSERM, Collège de France and Université de Paris (MT and SV).

We thank Prof. Ursula Jakob for providing the NemR^C106^ plasmid; Prof. Haike Antelman for providing the HypR plasmid; Daria Ezeriņa for several fruitful discussions; the beamline scientists at the Proxima 2 the beamline of the Soleil synchrotron facility and the beamline scientist Pierre Legrand of the Proxima 1 beamline at the Soleil Synchrotron facility for his help with data processing.

## Author contributions

A.I.K. developed architecture and design of the working version of Hypocrates biosensor, performed the in vitro experiments, analyzed and combined data, wrote the manuscript;

M-A.T. crystallized, collected data, and solved the structure of HypocratesCS, performed the in vitro circular dichroism experiments, pKa determination, fluorescence selectivity and sensitivity experiments, pre-steady state kinetic measurements of Hypocrates and HypR/NemR and wrote the manuscript;

A.S.P. performed experiments in eukaryotic cell culture;

M.T performed experiments in zebrafish;

K.W. helped with the crystal conditions optimization;

I.V.M. helped with X-ray structure refinement, and X-ray data deposition;

R.I.R. helped with the in vitro experiments (selectivity and sensitivity);

M.S.B. synthesized chemical compounds (NCT, ONOO-);

S.V. supervised the in vivo work;

J.M. supervised the in vitro and structural work and wrote the manuscript;

D.S.B. and V.V.B. created the general concept of the project, supervised the work of the project at all stages, wrote the manuscript.

## Methods

### Expression and purification of *S. aureus* HypR

Protein expression and purification were performed as described by Van Loi *et al.*^1^ with minor modifications. Briefly, the harvested cells were lysed using a Sonic VibraCell sonicator for 10 min, with 30 s sound/30 s pause with 61% amplitude. Cell debris was removed by centrifugation (45 min at 18,000 rpm, at 4°C; Avanti^®^ J-26xp centrifuge (BECKMAN COULTER^®^)), and the supernatant was in-batch incubated with Ni^2+^-Sepharose 6 Fast Flow beads (Cytiva) equilibrated with the binding buffer (20 mM HEPES/NaOH pH 7.5, 0.5 M NaCl and 10 mM imidazole) for 1 h at 4°C. The beads were then packed in a column coupled to an AKTA™ Pure system (GE Healthcare, Life Sciences). HypR was eluted using a linear gradient with elution buffer: 20 mM HEPES pH 7.5, 0.5 M NaCl and 0 to 500 mM (0-100%) imidazole. Protein purity was assessed on a nonreducing SDS-PAGE gel, and the pure fractions were collected, dialyzed (~20 mL sample/2 L dialysis buffer) overnight at 4°C against 20 mM HEPES pH 7.5 and 250 mM NaCl, and stored at −80°C in 20% glycerol.

### Expression and purification of *E. coli* NemR^C106^

The pET-21b(+)-NemR^C106^ plasmid ^2^, which contains only one cysteine (Cys106), was transformed in *E. coli* BL21 (DE3) cells. Cells were grown in Lysogeny Broth (LB) supplemented with 50 μg/mL of kanamycin at 37°C until the A_600_ reached 0.8. Isopropyl β-d-1-thiogalactopyranoside (IPTG) (0.5 mM) was used for the expression induction, followed by 3 h of incubation at 37°C. Harvested cells were then pelleted at 4°C, 5000 rpm for 15 min using the Avanti^®^ J-26xp centrifuge (Beckman Coulter^®^) and resuspended in lysis buffer composed of 50 mM Tris/HCl pH 8, 0.2 M NaCl, 1 mM DTT, 0.1 mg/ml 4-(2-aminoethyl) benzenesulfonyl fluoride hydrochloride (AEBSF), 1 μg/ml Leupeptin, 50 μg/ml Dnase I, and 20 mM MgCl_2_. Cells were disrupted and centrifuged as mentioned above. The supernatant was in-batch incubated with Ni^2+^-Sepharose 6 Fast Flow beads (Cytiva) equilibrated with 50 mM Tris/HCl pH 8, 0.2 M NaCl and 1 mM DTT for 1 h at 4°C. The beads were packed in a column, and the AKTA™ Pure system (GE Healthcare, Life Sciences) was used for purification. NemR^C106^ was eluted using a linear gradient with elution buffer consisting of 50 mM Tris/HCl pH 8, 0.2 M NaCl, 1 mM DTT and 0 to 700 mM (0-100%) imidazole. Following purification, protein purity was assessed on a nonreducing SDS-PAGE gel, and the pure fractions were dialyzed (~20 mL sample/2 L dialysis buffer) overnight at 4°C against the binding buffer and stored at −20°C.

### Molecular cloning procedures

Tersus Plus PCR Kit (Evrogen) was used for all amplification procedures. Primers are listed in **Supplementary Table 3**. An overlap extension PCR protocol was implemented to engineer NemR-cpYFP versions. Each reaction mix included NemR^C106^ N- and C-terminal fragments and cpYFP fragment in equal molar amounts. The DNA concentration was estimated with horizontal DNA electrophoresis in an agarose gel. The pQE30-HyPer-2 plasmid ^3^ was used as a template to amplify the cpYFP part. Two versions of this fragment (with SAG/G and SAG/GT linker pairs) were generated with the use of №1/№18 and №2/№19 primer pairs, respectively. The pET-21b(+)-NemR^C106^ plasmid was used as a template to amplify NemR^C106^ N- and C-terminal parts. All NemR^C106^ N-terminal parts were generated with the use of primer №3 and one of the primers from the №21-32 subset. All NemR^C106^ C-terminal parts were generated with the use of primer №20 and one of the primers from the №4-15 subset. Upon completion of the overlap extension PCR protocol, the target product was separated from the nontarget byproducts with horizontal DNA electrophoresis in agarose gel and purified with Cleanup Standard Kit (Evrogen). To engineer pQE30-NemR-cpYFP plasmids, the purified NemR-cpYFP constructs and intact pQE30 vector were incubated with BamHI and HindIII FastDigest™ enzymes in the corresponding buffer (Thermo Scientific) at 37°C for 20 minutes. The restricted polynucleotides were purified with Cleanup Standard Kit (Evrogen) and ligated with T4 DNA ligase in the corresponding buffer (Evrogen) at 14°C overnight. The molar vector/insert ratio was approximately 1:3 in all cases. The DNA concentration was estimated with horizontal DNA electrophoresis in an agarose gel. After incubation, the samples were transformed into *E. coli* XL1Blue cells, which were grown on LB-agar plates containing 100 μg/ml ampicillin for 14 h at 37°C. To detect colonies bearing the target plasmid, ScreenMix Kit (Evrogen) was used according to the manufacturer’s protocol. The positive colonies were then transferred to 100 μg/ml ampicillin LB and grown for 14 h at 37°C, 200 rpm (New Brunswick™ Excella^®^ E25). The resulting NemR-cpYFP-bearing vectors were purified with the use of Plasmid Miniprep Kit (Evrogen) according to the manufacturer’s protocol. The DNA concentration in the pure samples was measured with the use of a NanoDrop 2000 spectrophotometer (Thermo Scientific). The lack of any undesired mutations in the engineered genes was established by DNA sequencing (Evrogen).

An overlap extension PCR protocol was implemented to engineer inactivated HypocratesCS version (the first-generation control). The reaction mix included Hypocrates N- and C-terminal fragments with the desired substitution in equal molar amounts. The DNA concentration was estimated with horizontal DNA electrophoresis in an agarose gel. The pQE30-Hypocrates plasmid was used as a template to amplify both parts. The N- and C-terminal parts were generated with the use of №3/№33 and №16/№20 primer pairs, respectively. The reaction mix after overlap extension PCR was subjected to the same procedures as described above.

To transfer any NemR-cpYFP version from the pQE30 vector to the PCS2+ vector, the corresponding gene was amplified with the use of the №17/№34 primer pair and purified with Cleanup Standard Kit (Evrogen). The obtained construct and intact PCS2+ vector were incubated with ClaI and XbaI FastDigest™ enzymes in the corresponding buffer (Thermo Scientific) at 37°C for 20 minutes. The restricted polynucleotides were then subjected to the same procedures as described above.

### Functionality tests of NemR-cpYFP variants in *E. coli* cells

To obtain bacterial cells that express any of the NemR-cpYFP variants, the pQE30 vector bearing the desired gene was transformed to *E. coli* XL1Blue cells, after which they were grown on LB-agar plates containing 100 μg/ml ampicillin for 14 h at 37°C. In all cases, the bacterial density was controlled to achieve conditions in which the individual colonies were located at a distance of 1-2 mm from each other, as this parameter significantly affects the maturation and the redox state of the sensors. The fluorescence intensity of the cells was estimated with the use of an Olympus US SZX12 fluorescent binocular microscope. On the first day, all NemR-cpYFP versions were characterized by weak fluorescence, which was attributed to the fact that circularly permuted fluorescent proteins have destabilized structure and require more time for efficient maturation. Given that, the LB-agar plates were additionally incubated for 24 h at 17-20°C, as it is known that the maturation of circularly permuted fluorescent proteins proceeds better at lower temperatures.

To test the functionality of NemR-cpYFP variants, the bacterial biomass was transferred to 1 ml of PBS (137 mM NaCl, 2.7 mM KCl, 10 mM Na_2_HPO_4_, 1.8 mM KH_2_PO_4_, pH = 7.4) and resuspended with an automatic pipette. The fluorescence spectra (λ_ex_ = 425 nm or 500 nm) and the excitation spectra (λ_em_ = 525 nm) were recorded with the use of a Varian Cary Eclipse Fluorescence Spectrophotometer. The suspensions were treated with NaOCl aliquots to achieve the final oxidant concentration of 80 μM, after which the spectral measurements were repeated. In all cases, the samples were mixed by pipetting prior to the final spectra registration until the signal stabilization was observed. The data were analyzed with OriginPro 9.0 (OriginLab).

### Expression and purification of NemR-cpYFP variants, EYFP, intact cpYFP, HyPer-2 and SypHer3s

In the current work, two different protocols for Hypocrates expression and purification were used. Both of them led to obtaining the functional biosensor. Therefore, they should be considered to be equal.

#### Protocol 1

Xl1Blue cells were transformed with pQE30-Hypocrates plasmid, after which they were plated (LB-agar medium, 100 μg/ml ampicillin) and incubated for 14 h at 37°C. The bacterial density was controlled as described above. To achieve better protein folding and maturation, the plates were additionally incubated for 24 h at 17-20°C. Next, the cells were washed from the agar surface by ice-cold PBS, and the final volume of the suspension was adjusted to 24 ml with the same buffer. The number of plates used for a single purification procedure was twenty. The cells were destructed with the use of a Sonic VibraCell instrument in an ice bath (5 s sonication + 10 s pause cycle; total sonication time – 9 min; the amplitude – 32%). The obtained lysates were centrifuged for 20 min at 21,000 g and 4°C (Centrifuge 5424 R, Eppendorf) to precipitate insoluble fractions. The supernatants were collected and applied to a column filled with 5 ml of TALON Metal Affinity resin (Takara) previously equilibrated with ice-cold PBS. The column was washed with 50 ml of the same buffer to get rid of nontarget proteins. The elution step was performed by the addition of 10 ml of ice-cold PBS containing 250 mM imidazole, and the fraction with the target protein was collected on the basis of its bright yellow color. The elimination of imidazole was achieved by gel filtration on columns filled with 10 ml of Sephadex G-25 (GE Healthcare Life Sciences) previously equilibrated with ice-cold PBS. The pure protein sample was stored at 4°C for no more than 3 days. The addition of any reducing agents (such as β-mercaptoethanol) did not alter the properties of the protein – the sensor was obtained in its fully reduced form, even in their absence. Hypocrates samples, purified according to this protocol, were implemented for the following tests: the measurements of optical parameters, fluorescence spectra stability and reversibility experiments, fluorescence selectivity experiments, and measurements of MPO activity. Other primary NemR-cpYFP versions, EYFP, cpYFP, and SypHer3s were purified according to the same protocol as well as HyPer-2. However, in the case of the latter, all buffers, except for those used at the gel filtration step, contained 5 mM β-mercaptoethanol to avoid the oxidation of the sensor. The protein concentration in the final samples was measured with the use of Bicinchoninic Acid Kit for Protein Determination (Sigma-Aldrich) and a 96-well plate analyzer (Tecan Infinite 200 PRO).

#### Protocol 2

Shuffle^®^ T7 or XL1Blue cells were transformed with pQE30-Hypocrates plasmid, respectively. The cells were plated on LB-agar-ampicillin and incubated overnight at 37°C (for XL1Blue) and 30°C (for Shuffle^®^ T7). Plates were transferred to a 25°C incubator until they expressed the protein, as indicated by yellow colored colonies. At the next step, several colonies were transferred to 3 L of LB medium supplemented with 100 μg/mL ampicillin and incubated for 36 h at 25°C by rotating at 180 rpm. Cells were harvested, and the pellet was resuspended in lysis buffer composed of 40 mM Tris pH 7.5, 150 mM KCl, 10 mM MgSO_4_, 5 mM DTT, 0.1 mg/ml AEBSF, 1 μg/ml Leupeptin, 50 μg/ml DnaseI, and 20 mM MgCl_2_. Cells were lysed and centrifuged as performed for NemR^C106^, and the supernatant was in-batch incubated with Ni^2+^-Sepharose beads (Thermo Scientific) equilibrated with binding buffer: 40 mM Tris pH 7.5, 150 mM KCl, 10 mM MgSO_4_ and 1 mM DTT for 1 h at 4°C. After column packing, the AKTA™ Pure system (GE Healthcare, Life Sciences) was used to elute the protein using a binding buffer with 400 mM imidazole followed by size exclusion chromatography on a Superdex75 16/600 (GE Healthcare) column equilibrated in binding buffer. The purity of the protein was assessed on a nonreducing SDS-PAGE gel, and the pure fractions were collected and stored at −20°C. Hypocrates sample, purified according to this protocol, was implemented for the following tests: circular dichroism experiments, pKa determination, fluorescence selectivity experiments, fluorescence sensitivity experiments, presteady state kinetic measurements, HypocratesCS crystallization.

### N-chlorotaurine and NaONOO preparation

The preparation of N-chlorotaurine was carried out according to Patent DE4041703A (https://patents.google.com/patent/DE4041703A1/en). Chloramine T trihydrate (6.0 g, 21.3 mmol) was dissolved in dry methanol (50 mL). Finely powdered taurine (2.5 g, 20 mmol) was added, and the mixture was stirred for 20 h at room temperature (20-25°C). The solvent was removed on a rotary evaporator, and the residue was washed with isopropyl alcohol (3 times, 10 ml) and diethyl ether (3 times, 35 ml). The white solid was dried in vacuum (5 mmHg, 1 h). The NMR analysis of the product (DMSO-d6) showed the absence of aromatic protons. The product was stored at −20°C.

The preparation of NaONOO solution was carried out according to Uppu ^4^. NaOH (4.0 g, 0.10 mol) was dissolved in water (35 mL). The mixture was cooled in an ice bath to 5-0°C, and a solution of 35% H_2_O_2_ (11 ml, approximately 0.11 mol) and EDTA (solid, 75 mg) were added. Liquid isoamyl nitrite (13.5 ml, 0.10 mol) was added, and the mixture was vigorously stirred at room temperature (~25°C) for 5 h. The mixture was diluted with dichloromethane (100 ml), and the water phase was separated and washed additionally with dichloromethane (5 times, 100 ml each). The unreacted H_2_O_2_ was then removed by passing the aqueous phase through manganese dioxide (10-15 g, 5 mm layer). The resulting solution was additionally filtered from traces of MnO_2_, and the traces of dichloromethane were removed in vacuum (5 mmHg, 1 h). The resulting NaONOO solution was used in the further experiments. The solution was stored at −20°C. The concentration of ONOO^−^ ions was determined before each usage using spectrophotometry (Varian Cary 5000 Spectrophotometer). For these measurements, the solution was diluted with NaOH solution (pure water, 0.1 M concentration). Concentration was determined using Lambert-Beer’s law: ∊ at 302 nm = 1670 M^−1^cm^−1^ for ONOO^−^ ions.

### Measurements of the optical parameters of NemR-cpYFP variants

To measure the brightness of the primary NemR-cpYFP versions, the proteins were diluted in PBS to equimolar concentrations (according to Bicinchoninic Acid Kit). Purified EYFP served as the comparison control. The absorbance and fluorescence excitation spectra (λ_em_ = 513 nm and 533 nm for NemR-cpYFP variants and EYFP, respectively) of the samples were recorded with the use of a Varian Cary 5000 Spectrophotometer or a Varian Cary Eclipse Fluorescence Spectrophotometer. The molar extinction coefficients (ε) were calculated according to the following equation –ε = A/(C•L), where A was the optical density at the studied absorption maximum, C was the protein concentration (M), and L was the optical path length (cm). The fluorescence quantum yields (QY) were calculated according to the following equation – QY_NemR-cpYFP_ = QY_EYFP_•(A_EYFP_•Em_NemR-cpYFP_/(A_NemR-cpYFP_•Em_EYFP_)), where A was the optical density at the studied absorption maximum, and Em was the emission intensity at the studied excitation maximum (λ_ex_ = 425 nm or 500 nm for NemR-cpYFP variants, and 519 nm for EYFP). QY_EYFP_ is a standard value of 0.67 according to the literature (Fpbase ID: 8DNLG). The data were analyzed with OriginPro 9.0 (OriginLab).

The purified sensor samples might contain not fully folded and matured molecules, reducing the accuracy of the optical parameters’ measurements. Therefore, the molar extinction coefficients and Qys of Hypocrates were investigated in more detail. To estimate the concentration of fully matured chromophores, the samples of Hypocrates and EYFP were mixed with 0.1 M NaOH at the volume ratio of 1:1 and incubated for 5 minutes. In the described conditions, yellow fluorescent proteins undergo denaturation, and mature chromophores are converted to the form absorbing at 445 nm with ε = 44000 M^−1^cm^−1^ ^5^. To investigate how reducing and oxidizing agents alter the optical parameters, some of the Hypocrates samples were incubated in the presence of 0.5 mM N-chlorotaurine or 5 mM DTT for 30 minutes prior to spectra registration. All of the following procedures were carried out as described above. The data were analyzed with OriginPro 9.0 (OriginLab).

### Fluorescence spectra stability and reversibility experiments

To investigate whether high oxidant concentrations damage the proteins, purified Hypocrates samples (0.5 μM) were treated with saturating oxidant concentrations (5-10 μM), and their fluorescence excitation spectra (λ_em_ = 520 nm) were recorded. Next, the aliquots of corresponding oxidants were added to achieve extremely high concentrations (100 μM), and the same measurements were carried out. In all cases, the samples were mixed by pipetting prior to the final spectra registration until the signal stabilization was observed. NaOCl and N-chlorotaurine were tested in PBS, while NaONOO was tested in sodium phosphate buffer to avoid possible OCl^−^ generation in the system. In the last case, the protein aliquots were transferred to the corresponding buffer with the use of Amicon Ultra-0.5 Centrifugal Filter Units (Millipore). The NaOCl sensitivities of intact cpYFP, SypHer3s and HyPer-2 purified proteins were investigated according to the same protocol. The measurements were performed with the use of a Varian Cary Eclipse Fluorescence Spectrophotometer. The data were analyzed with OriginPro 9.0 (OriginLab).

To investigate whether Hypocrates oxidation is reversible, purified protein samples (0.5-2 μM) were treated with saturating oxidant concentrations (5-50 μM) and incubated for 5 min, after which the fluorescence excitation spectra were recorded. Next, DTT was added to the reaction mix to the final concentration of 1-5 mM, and the probes were incubated for 40-60 min prior to the spectra registration. In some cases, two additional control probes (intact and with the same oxidant concentration) were prepared and incubated for the same time to control for possible artifacts caused by prolonged atmosphere exposure. The measurements were performed with the use of either a Varian Cary Eclipse Fluorescence Spectrophotometer or an LS55 luminescence spectrophotometer. The data were analyzed with OriginPro 9.0 (OriginLab).

### Biosensor secondary structural changes with circular dichroism

Changes in the overall secondary structure of the Hypocrates between its reduced and oxidized (NaOCl or H_2_O_2_) forms were evaluated with circular dichroism (CD) spectroscopy. The protein was reduced with 30 mM DTT for 30 min at room temperature. A Hi-Trap® desalting column (GE Healthcare), equilibrated with 20 mM sodium phosphate buffer pH 7.4, was used to remove excess DTT. To prepare the oxidized samples, Hypocrates (25 μM) was incubated for 10 min at room temperature with different concentrations of NaOCl (1:1, 1:5 and 1:10 ratios) or H_2_O_2_ (1:1, 1:3 and 1:6 ratios of protein to oxidant concentration), with a reaction buffer composed of 20 mM sodium phosphate, pH 7.4, and 200 mM sodium fluoride. Micro Bio-Spin^®^ Chromatography Columns (BIO-RAD), equilibrated with the same buffer, were used to remove the oxidants. Following sample preparation, a Jasco J-810 spectropolarimeter was used to analyze 4 μM of each sample at 25°C in a quartz cuvette with a 1-mm path length. Far-UV CD spectra (190-260 nm) were measured, and the data were analyzed with GraphPad Prism8 and OriginPro 9.0 (OriginLab).

To determine whether the overall secondary structure could be restored, DTT was used. NaOCl-oxidized Hypocrates (1:10 protein/oxidant ratio) was incubated with 200 μM DTT for 5 min at room temperature. The background from the buffer and the addition of 200 μM DTT was subtracted.

### pKa determination of reduced and oxidized NemR-cpYFP versions

To determine the pKa of Hypocrates, the protein was reduced with DTT and buffer-exchanged into 100 mM sodium phosphate buffer using a Hi-Trap® desalting column (GE Healthcare). Reduced Hypocrates (0.5 μM) in the presence or absence of 12.5 μM oxidants (NaOCl and NCT) was diluted in a polybuffer solution with several pH values (0.5 pH unit intervals), and the excitation spectra (with λ_em_ = 555 nm) were recorded after 5 min of incubation at 25°C using a SpectraMax iD5 plate reader (Molecular Devices). The polybuffer solution consisted of sodium acetate (10 mM), sodium phosphate (10 mM), sodium borate (10 mM) and sodium citrate (10 mM). The Ex_500_/Ex_417_ ratio was plotted as a function of increasing buffer pH. For each measurement, at least three independent replicates were performed, and the data were analyzed using GraphPad Prism8 and OriginPro 9.0 (OriginLab). The pKa of reduced HypocratesCS was determined as described for reduced Hypocrates.

### Fluorescence selectivity experiments

Aliquots of the sensor (2 μM) were incubated with different oxidants: NaOCl (50 μM), N-chlorotaurine (50 μM), H_2_O_2_ (50 μM), glutathione (GSH; 500 μM), glutathione disulfide (GSSG; 500 μM), MAHMA NONOate (NO^•^ generator; 50 μM), Na_2_S (50 μM), NaONOO (50 μM), and X/XOX system (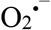 generator; 50 μM + 0.05 U/ml). Both NaONOO and X/XOX samples contained catalase (0.1 μM) as an additional control, which removed H_2_O_2_ generated during the reaction. The samples were incubated for 5 min at 25°C, and the ratiometric fluorescence changes were monitored by an excitation scan (λ_em_ = 515 nm) using an LS55 luminescence spectrophotometer (PerkinElmer). For each concentration, at least three independent experimental measurements were performed. The changes in intrinsic Trp fluorescence were measured using an emission scan (λ_ex_ = 295 nm). The data were analyzed with OriginPro 9.0 (OriginLab).

### Fluorescence sensitivity experiments

Hypocrates sensitivity experiments were performed in 100 mM sodium phosphate buffer. Aliquots of the purified protein (0.5 μM) were incubated with increasing concentrations of NaOCl and NCT for 5 min at 25°C. The excitation scans (with λ_em_= 555 nm) were recorded with the use of a SpectraMax iD5 plate reader (Molecular Devices). The Ex_500_/Ex_417_ ratio, which represents the ratio between fluorescence excited at 500 nm and at 417 nm, was plotted as a function of increasing oxidant concentration. The initial linear part of the hyperbolic curve was analyzed using linear regression, where the slope values represent the sensitivity towards the corresponding oxidants. For each measurement, at least two independent replicates were performed, and the data were analyzed using GraphPad Prism8 and OriginPro 9.0 (OriginLab).

### Presteady-state kinetic measurements

Presteady-state kinetic measurements were performed using a stopped-flow apparatus coupled to a fluorescence detector (Applied Photophysics SV20). For HypR, changes in intrinsic Tyr fluorescence were measured using a >305 nm cut-off filter (λ_ex_ = 274 nm). For NemR^C106^, changes in intrinsic Trp fluorescence were measured using a >320 nm cut-off filter (λ_ex_ = 295 nm). For Hypocrates, changes in the cpYFP chromophore fluorescence were measured using a >515 nm cut-off filter (λ_ex_ = 485 nm). The excitation and emission wavelengths for each protein were determined prior to the stopped-flow experiments using an LS55 luminescence spectrophotometer (PerkinElmer). The oxidant concentration range required to lead to fluorescence changes for each protein was also determined.

Prior to the experiment, the samples were reduced with 30 mM DTT for 30 min at room temperature. A Hi-Trap® desalting column (GE Healthcare), equilibrated with argon-flushed 100 mM sodium phosphate pH 7.4 buffer, was used to remove excess DTT. To determine the second-order rate constants, 0.5 μM Hypocrates or 1 μM NemR or HypR was mixed with increasing concentrations of an oxidant (NaOCl or N-chlorotaurine) in a reaction medium of 100 mM sodium phosphate buffer at 25°C. Changes in fluorescence were monitored, and the obtained curves were fitted with a single exponential equation (Y = A.e^(−*k*obs.t)^ + offset). For each oxidant concentration, the observed rate constant (*k*_obs_) was determined. The *k*_obs_ values were plotted against the different oxidant concentrations, and linear regression was used to obtain the second-order rate constants from the slope (GraphPad Prism8 and OriginPro 9.0). For each concentration, at least two independent experimental measurements were performed.

### HypocratesCS crystallization, X-ray data collection, and structure determination

HypocratesCS was crystallized at 7 mg/mL concentration at 10°C or 20°C using the hanging-drop vapor diffusion method with Tris (0.1 M, pH 8), CaCl_2_ (0.1 M), MgCl_2_ (0.1 M) and PE15/4 (15%) as a precipitant solution. The drops were composed of 1 μL of protein and 1 μL of precipitant solution. To obtain larger and better diffracting crystals, the small needles obtained within the above crystallization condition were used for microseeding.

For X-ray data collection, the cryo-protectant used was the same as the precipitant solution, but with 30% PE15/4. X-ray data were collected at 100 K at the Proxima 2 beamline of the Soleil synchrotron facility, at a wavelength of 0.980113 Å, and processed using XDS ^6^. The STARANISO server was used to perform anisotropic correction of the data ^7^. The structure of HypocratesCS was solved by molecular replacement using Phase ^8^ from the Phenix suite ^9^, using both the *E. coli* NemR (PDB: 4YZE) (100% identical to the sensory domain of HypocratesCS) and the cpYFP-based calcium biosensor (PDB: 3O77) (98% identity to the cpYFP) as search models. Coot ^10^ was used to manually complete the building of the structure, and the refinement was done using Phenix. Refine ^11^ from the Phenix suite. Analysis of the Ramachandran plot showed that 96.59% of the residues are in the most favored areas of the Ramachandran plot, 2.93% in additionally allowed areas, and 0.49% in disallowed areas. The data collection statistics and refinement parameters are summarized in **Supplementary Table 2**.

### Measurements of MPO activity with purified Hypocrates

To test whether Hypocrates is capable of visualizing MPO activity *in vitro*, the purified sensor was incubated with 0.1 U/ml human MPO and 100 μM H_2_O_2_ for 10 min in PBS, after which the fluorescence excitation spectrum (λ_em_ = 525 nm) was recorded with the use of a Varian Cary Eclipse Fluorescence Spectrophotometer. The probes that contained only one component of the MPO-H_2_O_2_ system were treated according to the same protocol to control for nonspecific fluorescence changes. The data were analyzed using OriginPro 9.0 (OriginLab).

To record the dynamics of HOCl production by MPO *in vitro*, purified Hypocrates was incubated in the presence of 0.1 U/ml human MPO, and the intensities of fluorescence (λ_em_ = 525 nm) excited at 425 nm and 500 nm were collected each 2.4 s with the use of a Varian Cary Eclipse Fluorescence Spectrophotometer. To start the MPO reaction, 100 μM H_2_O_2_ was added to the reaction mix. A probe without MPO was treated according to the same protocol to control for H_2_O_2_-induced fluorescence changes. A probe without MPO and without H_2_O_2_ addition was treated according to the same protocol to control for fluorescence changes attributed to prolonged incubation. For all three samples, the Ex_500_/Ex_425_ ratio was calculated as a function of time, and the first two curves were normalized by the third one. The data were analyzed using OriginPro 9.0 (OriginLab).

### NaOCl visualization with Hypocrates, HypocratesCS and SypHer3s in HeLa Kyoto cells

HeLa Kyoto cells were cultured in DMEM (PanEko) supplemented with 10% FBS (Biosera), 2 mM L-glutamine (PanEko), 50 units/ml penicillin (PanEko) and 50 μg/ml streptomycin (PanEko) at 37 °C in atmosphere containing 5% CO_2_. Сells were passaged every 2-3 days. For transfection, cells were seeded into 35-mm glass-bottom dishes (SPL Lifesciences). After 24 h, cells were transfected with the plasmid of the required sensor using FuGene HD transfection reagent (Promega) according to the manufacturer’s protocol. Fluorescent microscopy was performed on the next day after transfection with a Leica DMI 6000 microscope, equipped with an HCX PL Apo CS 40.0 × 1.25 Oil UV objective, CFP (excitation filter BP 436/20, dichromatic mirror 455, suppression filter BP 480/40) and GFP (excitation filter BP 470/40, dichromatic mirror 500, suppression filter BP 525/50) filter cubes. A 10 mM stock solution of NaOCl (EMPLURA) in Milli-Q water was freshly prepared before cell imaging. Cell culture medium was replaced with 900 μL of PBS, and baseline fluorescence was detected for several minutes. PBS was chosen as an inorganic imaging medium because NaOCl, being a strong oxidant, can react with the nitrogen-containing components of the medium and thus introduce inaccuracy to the results. The desired amount of NaOCl stock was diluted in 100 μL of PBS just before addition, and the final concentration of NaOCl in the sample was in the range of 10-40 μM. All measurements were taken at room temperature because the maximum response amplitude of Hypocrates is being reduced as a result of heating. The responses of sensors were calculated as ratios of fluorescence intensities excited at 500 nm and at 425 nm (Ex_500_/Ex_425_ ratio) and normalized to the signal of the probe on the first image of the series. Processing of the images and quantification of results were performed using Fiji (https://fiji.sc), Excel (Microsoft) and OriginPro 9.0 (OriginLab).

### *Danio rerio* tail fin inflammation model

For the tail fin amputation experiment, mRNAs of Hypocrates, HypocratesCS and HyPerRed were *in vitro* synthesized using mMessage mMachine Transcription Kit (Invitrogen) according to manufacturer’s manual. For transient expression of the biosensors in zebrafish larvae, 80 ng/μL of Hypocrates or HypocratesCS mRNA and 50 ng/μL of HyPerRed mRNA were co-injected into 1-cell-stage embryos. The zebrafish embryos were maintained in egg water containing 0.2 mM N-phenylthiourea (PTU; Sigma) to prevent pigment formation at 28 °C. Fluorescence imaging was performed 48 h postfertilization (hpf). Larvae were anesthetized in 0.02% MS-222, tricaine (Sigma), embedded in low-melting agarose (0.8%) and then subjected to tail fin amputation under a stereoscopic microscope. Imaging was performed with a CSU-W1 Yokogawa spinning disk coupled to a Zeiss Axio Observer Z1 inverted microscope equipped with a sCMOS Hamamatsu camera and a 25x (Zeiss 0.8 Imm WD: 0.19 mm) oil objective. DPSS 100 mW 405 nm and 150 mW 491 nm lasers and a 525/50 bandpass excitation filter were used for Hypocrates and HypocratesCS imaging. A 100 mW 561 nm laser and a 595/50 bandpass filter were used for HyPerRed imaging. To quantify the response, fluorescent signal at the amputation plane was normalized to the mean fluorescence of the tail before amputation. A statistical two-way ANOVA test with a Tukey’s multiple comparisons posttest was then performed.

