## Supplementary figures, tables for "Monitoring oxidative inflammatory processes in live cells and tissue with Hypocrates, a genetically encoded biosensor for hypochlorite": Hypocrates_Supplementary.pdf

### Supplementary Information

#### Supplementary figures

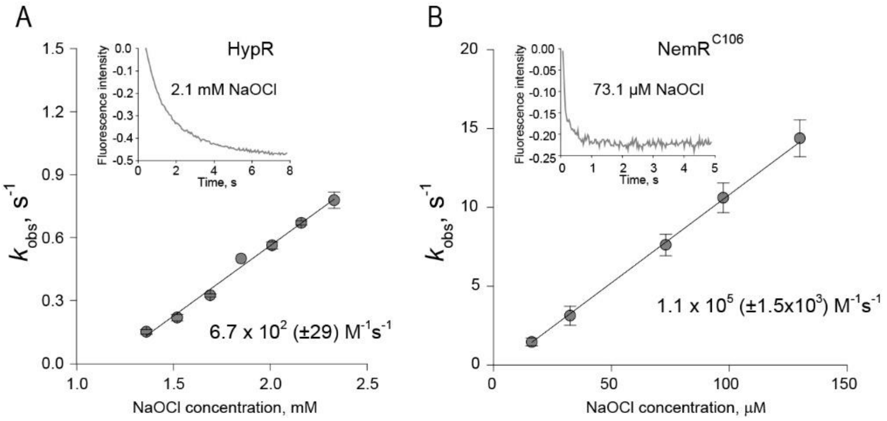


**Supplementary Figure 1**. $NemR^{C106}$ senses NaOCl 160-fold faster than HypR. The intrinsic Tyr (**A**) and Trp (**B**) fluorescence changes of HypR and $NemR^{C106}$ at increasing NaOCl concentration in function of time (inserts) are shown. Note, as HypR has no Trp in its sequence, we followed the Tyr fluorescence change. The curves were fitted to a single exponential to obtain the observed rate constants ($k_{obs}$), which were plotted as a function of increasing NaOCl concentration. From the slope, the second-order rate constant was obtained. The second-order rate constant of $NemR^{C106}$ is 1.1 x $10^5$ (± 1.5 x $10^3$) $M^{-1}s^{-1}$ and the one of HypR is 6.7 x $10^2$ (± 29) $M^{-1}s^{-1}$. The data are presented as the mean ± SD, n ≥ 2.

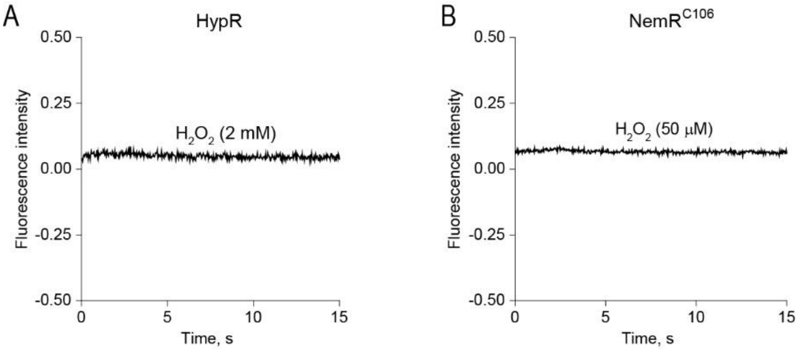


**Supplementary Figure 2**. HypR and $NemR^{C106}$ do not change their intrinsic fluorescence with $H_2O_2$.The changes of intrinsic fluorescence in function of time of HypR (**A**) and $NemR^{C106}$ (**B**) in the presence of $H_2O_2$ are shown. To confirm a higher specificity for NaOCl than for $H_2O_2$, the same concentration as for NaOCl was tested, 50 µM for $NemR^{C106}$ and 2 mM for HypR.

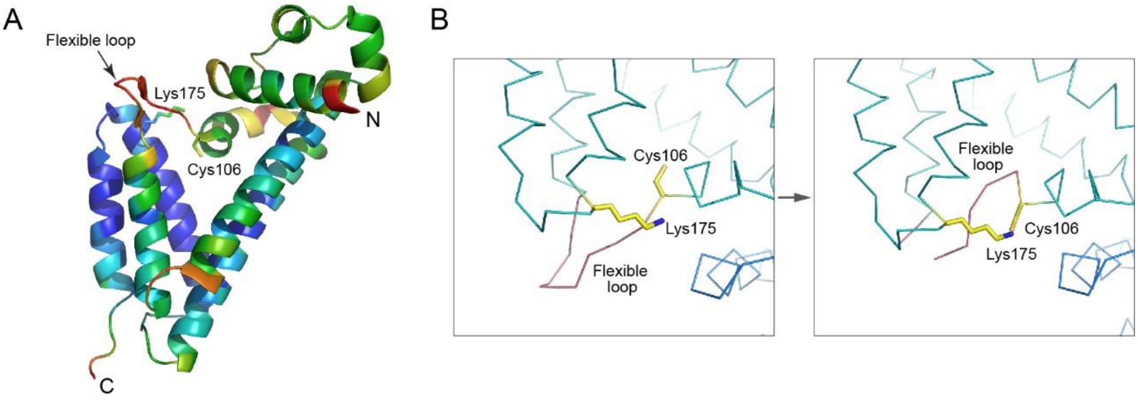


**Supplementary Figure 3**. The flexible regions of NemR$^{C106}$ (**A**) The structure of NemR$^{C106}$ (PDB ID: 4YZE) colored by b-factors (Cα) is shown. The color spectrum shows the flexibility going from most flexible (red) to most rigid (blue). (**B**) Different conformations of the Cys106 side chain can be found in the crystallographic asymmetrical unit. It has been suggested that $ClO^-$ induced oxidation leads to the transition from the left structure to the right one, affecting the flexible loop organization.

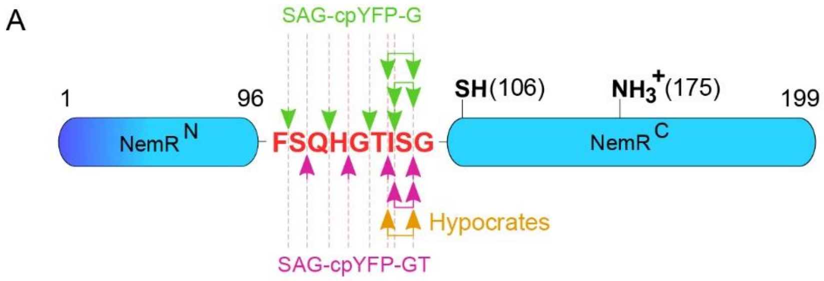


B

| Integration position | Max. response amplitude | Brightness at 425 nm (% of Hypocrates) | Brightness at 500 nm (% of Hypocrates) |
|---|---|---|---|
| 104-105 (SAG/GT) | ~ 1.2 | ~ 132 | ~ 82 |
| 103-105 (SAG/G) | ~ 1.2 | ~ 71 | ~ 52 |
| 103-105 (SAG/GT) | ~ 1.3 | ~ 143 | ~ 146 |
| 102-105 (SAG/G) | ~ 1.3 | ~ 78 | ~ 49 |
| 102-105 (SAG/GT) | ~ 1.6 | 100 | 100 |

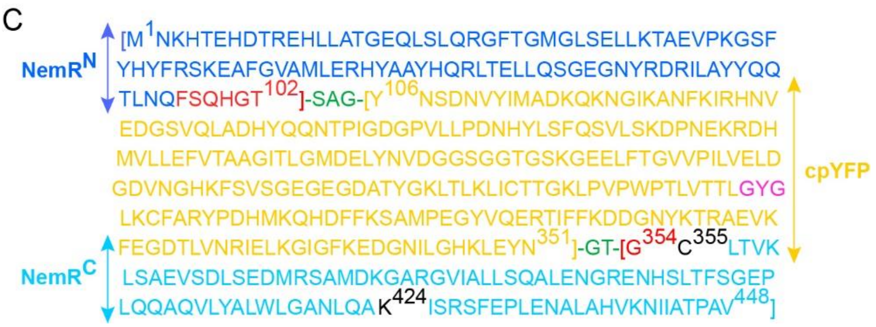


**Supplementary Figure 4**. (**A**) Different cpYFP inserting positions that were used for the development of the primary versions of NemR-cpYFP biosensor. The upper numbering on the scheme represent the amino acids corresponding to wild type NemR. Single and double arrows represent cpYFP insertions without and with deletions, respectively. (**B**) Characteristics of the selected primary versions of NemR-cpYFP biosensor (purified proteins). (**C**) Hypocrates primary structure. The color legend is the following: blue/cyan – NemR$^{C106}$ derived parts, yellow – cpYFP, red – the flexible loop, green – the linkers connecting NemR$^{C106}$ and cpYFP, black – the key Cys and Lys residues, pink – the chromophore triad.

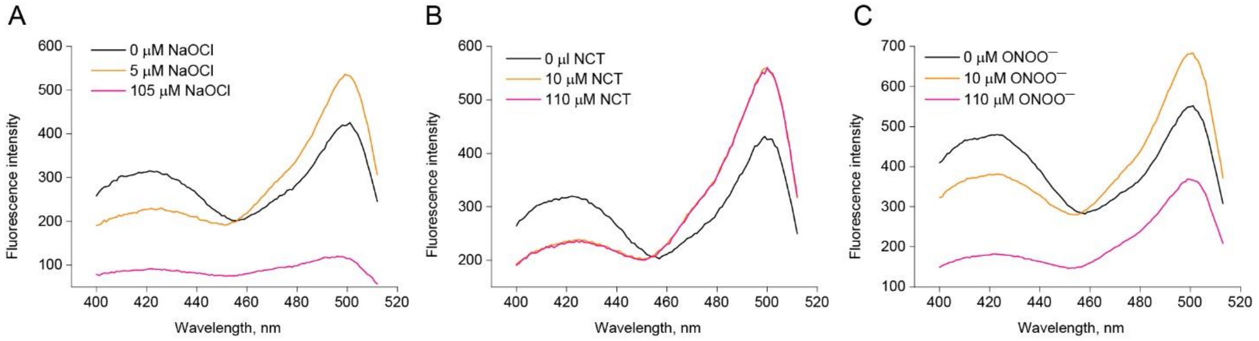


**Supplementary Figure 5**. (**A-C**) The degradation resistance of Hypocrates in the presence of high concentrations of oxidants: (**A**) NaOCl, (**B**) NCT, and (**C**) NaONOO. NCT does not induce fluorescence quenching apparently due to its lower reactivity and higher specificity toward sulfur containing amino acids. In all panels protein concentrations were 0.5 μM.

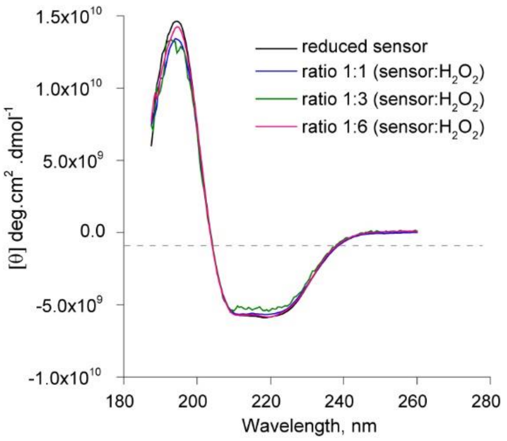


**Supplementary Figure 6**. Far-UV circular dichroism spectra of reduced and $H_2O_2$ oxidized: 1:1 ratio, 1:3 ratio and 1:6 ratio Hypocrates vs. oxidant, are shown. Addition of $H_2O_2$ does not induce change to the overall secondary structure of the biosensor.

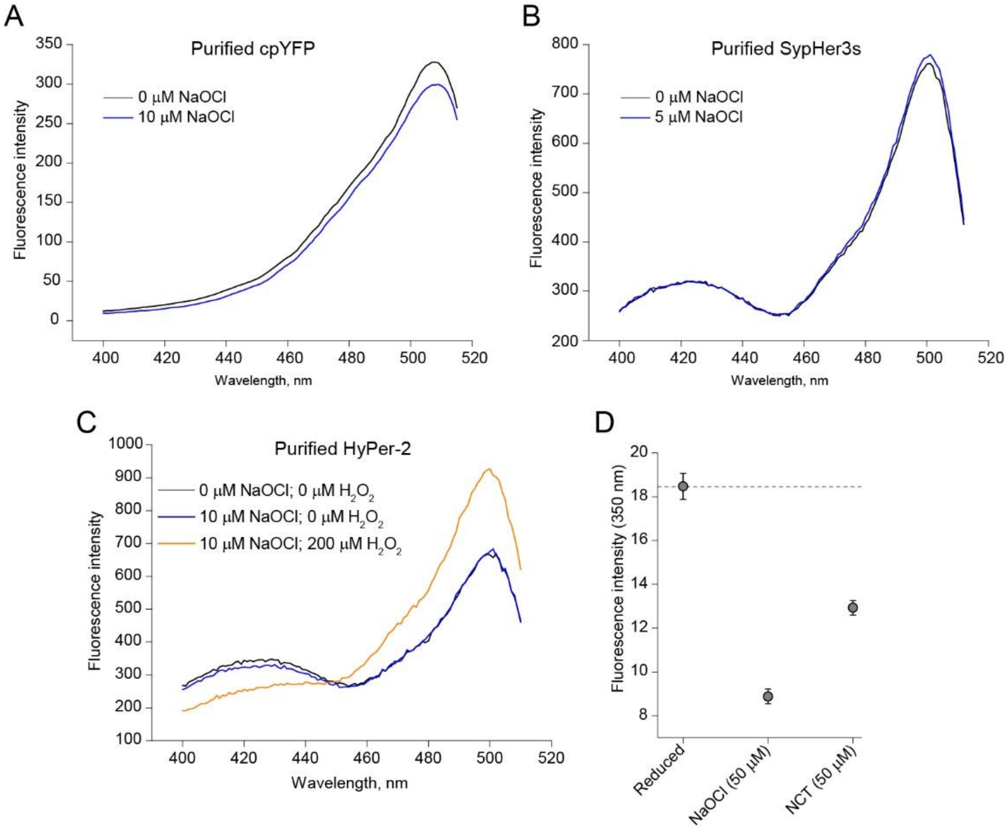


**Supplementary Figure 7. The response of different cpYFP-based probes to NaOCl.** (**A**) In the presence of NaOCl, the fluorescence excitation spectrum of purified cpYFP shows minor quenching due to apparent protein damaging. (**B**) The fluorescence excitation spectrum of the purified pH biosensor SypHer3s is resistant to NaOCl treatment. (**C**) Purified $H_2O_2$ biosensor HyPer2 does not react with NaOCl, while the addition of $H_2O_2$ induces a pronounced ratiometric response. (**D**) The intrinsic Trp fluorescence change of initial NemR$^{C106}$ in the presence of NaOCl and NCT. The data are presented as a mean ± SEM, n = 3. Protein concentration was 0.5 μM in panels **A-C** and 2 μM in panel **D**.

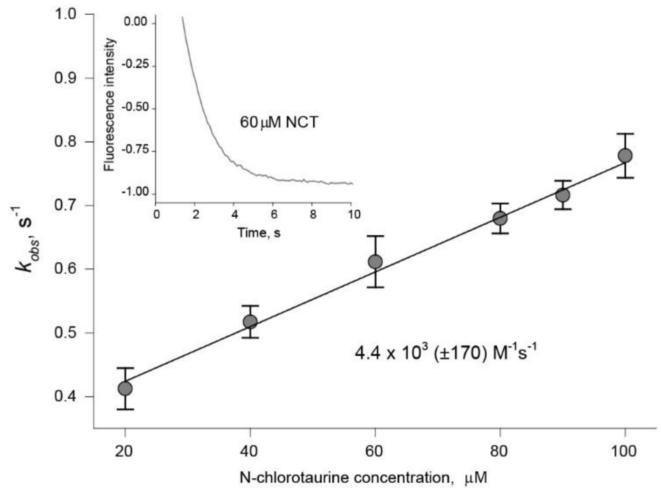


**Supplementary Figure 8.** The kinetics of NemR$^{C106}$ for N-chlorotaurine. Changes in intrinsic Trp fluorescence were measured in function of time (insert). The curves were fitted to a single exponential to obtain the observed rate constants ($k_{obs}$), which were plotted as a function of different NCT concentrations (20-100 μM). The second-order rate constant of 4.4 x $10^3$ (± 170 $M^{-1}s^{-1}$) was determined from the slope of the straight-line. The data are presented as a mean ± SD, n ≥ 2.

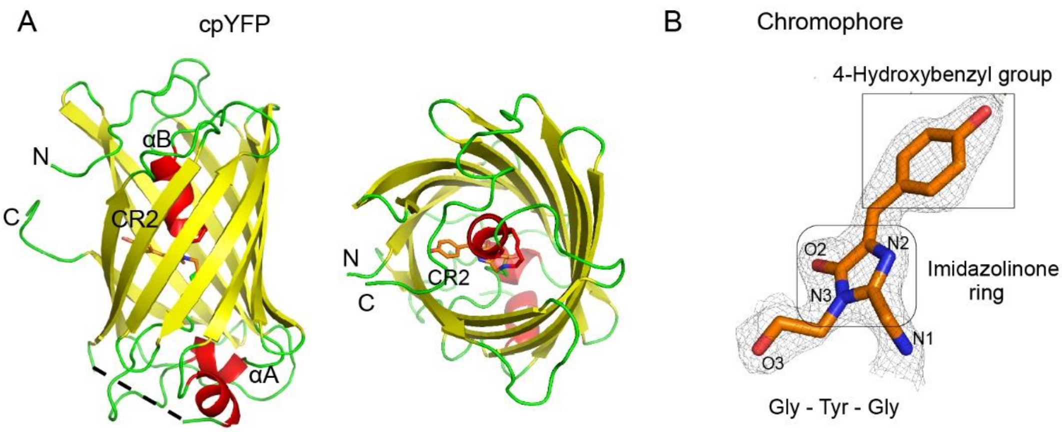


**Supplementary Figure 9.** The overall cpYFP architecture and the chromophore structure. The β-barrel of cpYFP (side and top views are shown) (**A**) is composed of 11 anti-parallel β-strands (yellow), which are connected via loops (green), and three α-helices (αA, αB and αC - red). The chromophore is located next to the C-terminal end of the αB-helix. Due to the high flexibility of an exposed loop (residues 191 to 207 - black dotted line), no defined electron density was observed for this region. (**B**) The chromophore structure is shown in orange and is modeled in cis configuration in the electron density map. The 4-hydroxybenzyl group and the imidazolinone ring are indicated by a black box.

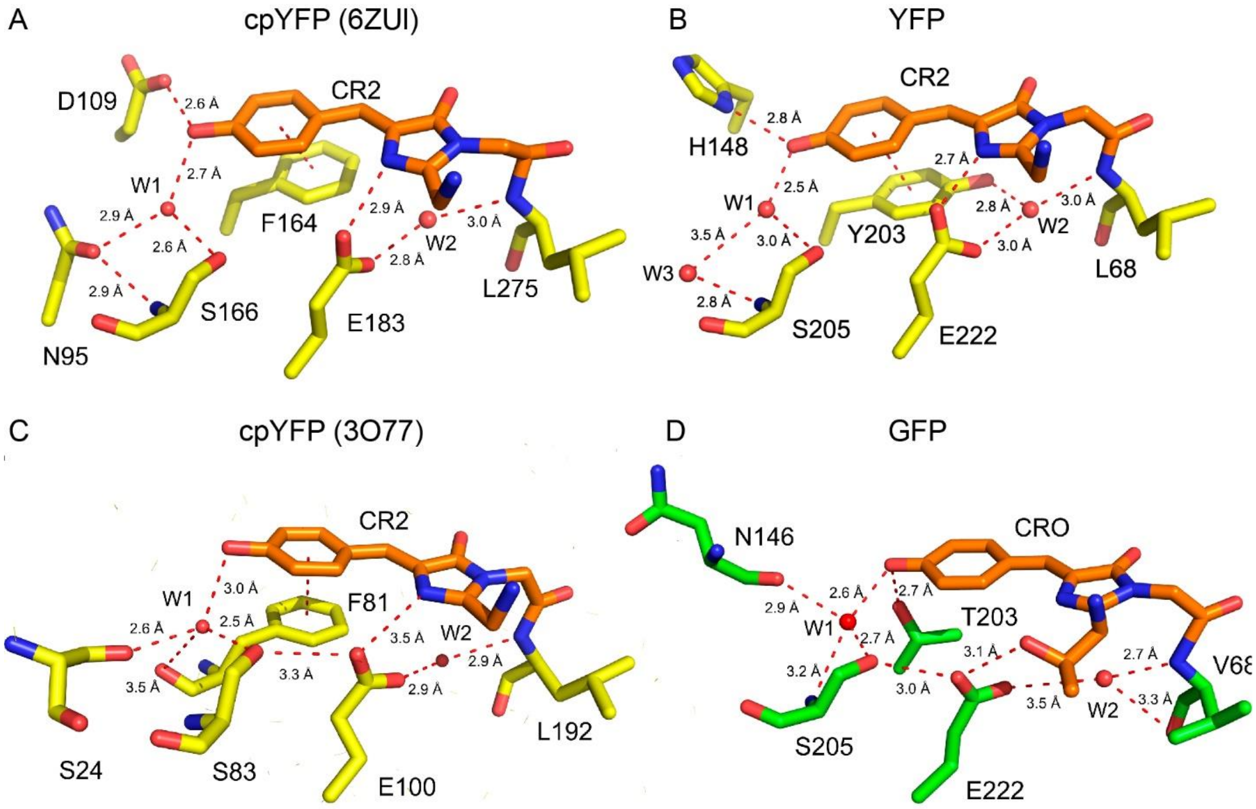


**Supplementary Figure 10.** The ESPT pathway is different for both cpYFPs, YFP, and GFP. (**A, B**) The CR2 environments of cpYFP (PDB ID: 6ZUI) and YFP (PDB ID: 1YFP) are shown. The positions of the two water molecules (W1 and W2) are conserved. In YFP, the CR2 oxygen is nearly in contact with bulk solvent through the two water molecules (W1 and W3) (panel B). In cpYFP of Hypocrates, the position of W3 is taken by OD1 of N95 (panel A), making the CR2 less solvent-exposed. In YFP, W2 has an extra H-bond with the Y203, which is absent in cpYFP. In cpYFP, the CR2 oxygen will be more deprotonated compared to the YFP because of H-bonding with OD2 of D109. In YFP, partial proton sharing with nitrogen ND1 of H148 will render this CR2 oxygen neutral. (**C**) The structural environment of the CR2 chromophore of the $Ca^{2+}$ sensor Case16 (PDB ID: 3O77) is shown. S24 links the sensing domain with the chromophore of cpYFP. (**D**) The structural environment of the CRO chromophore of GFP (PDB ID: 2B3P) is shown. T203 stabilizes a negative charge on the CRO oxygen.

**Supplementary Table 1. Optical properties of Hypocrates protein in fully reduced and fully oxidized states. Brightness is calculated as a product of extinction coefficient and quantum yield.**

| Wavelength, nm | Reduced | Oxidized |
|---|---|---|
| | Quantum Yield | |
| **425** | ~ 0.19 | ~ 0.15 |
| **500** | ~ 0.82 | ~ 0.83 |
| | Molar extinction coefficient, $M^{-1}cm^{-1}$ | |
| **425** | ~ 31200 | ~ 29600 |
| **500** | ~ 12000 | ~ 16700 |
| | Brightness | |
| **425** | ~ 5900 | ~ 4400 |
| **500** | ~ 9900 | ~ 13900 |

**Supplementary Table 2. X-ray data collection and refinement statistics**

| | HypocratesCS |
|---|---|
| **Data collection** | |
| **Space group** | C2221 |
| **Cell dimensions** | |
| ***a*, *b*, *c* (Å)** | 90.256, 95.463, 106.296 |
| **α, β, γ (°)** | 90.000, 90.000, 90.000 |
| **Resolution (Å)** | 47.72-2.10 (2.15-2.10)* |
| **$R_{merge}$ (%)** | 7.9 (62.5) |
| **$I$ / σ$I$** | 16.1 (2.1) |
| **Spherical completeness (%)** | 92.1 (68.3) |
| **Ellipsoidal completeness (%)** | 83.0 (46.6) |
| **Redundancy** | 9.3 (6.0) |
| **Refinement** | |
| **Resolution (Å)** | 47.72-2.10 |
| **No. reflections** | 22361 |
| **$R_{work}$ / $R_{free}$** | 20.12/26.75 |
| **No. atoms** | |
| **Protein** | 3284 |
| **Ligand/ion** | na |
| **Water** | 258 |
| ***B*-factors** | |
| **Protein** | 33.26 |
| **Ligand/ion** | na |
| **Water** | 22.69 |
| **R.m.s. deviations** | |
| **Bond lengths (Å)** | 0.007 |
| **Bond angles (°)** | 1.045 |

1 crystal was used to solve the HypocratesCS crystal structure

*Values in parentheses are for highest-resolution shell.

**Supplementary Table 3. The primers used in this work to engineer NemR-cpYFP versions**

"Ins." means "insertion position"

| Notification | № | Direct primer | № | Reverse primer |
|---|---|---|---|---|
| **SAG-cpYFP-G** | 1 | tctgcaggctacaacagcgacaacgtctat atcatggcc | 18 | accgttgtactccagcttgtgcccca |
| **SAG-cpYFP-GT** | 2 | tccgccggctacaacagcgacaacgtctat atcatggcc | 19 | ggtgccgttgtactccagcttgtgcccca |
| **NemR edges (pQE)** | 3 | atatatggatccatgaacaaacacaccga acatgatactcgc | 20 | Atatataagcttctaaacggcaggcgtcgca ataatgtttttac |
| **Ins. 97-98/G** | 4 | ggcacaagctggagtacaacggtagccaa catggaaccatcagtggttg | 21 | gttgtcgctgttgtagcctgcagaaaactggt tcagtgtttgctggtaataagcca |
| **Ins. 98-99/GT** | 5 | acaagctggagtacaacggcacccaacat ggaaccatcagtggttgcct | 22 | gctgttgtagccggcggagctaaactggttc agtgtttgctggtaataagc |
| **Ins. 99-100/G** | 6 | ggcacaagctggagtacaacggtcatgga accatcagtggttgcctgac | 23 | gttgtcgctgttgtagcctgcagattggctaa actggttcagtgtttgctggtaataag |
| **Ins. 100-101/GT** | 7 | acaagctggagtacaacggcaccggaacc atcagtggttgcctgacag | 24 | Gctgttgtagccggcggaatgttggctaaac tggttcagtgtttgctg |
| **Ins. 101-102/G** | 8 | ggcacaagctggagtacaacggtaccatc agtggttgcctgacagtaaaactc | 25 | gttgtcgctgttgtagcctgcagatccatgttg gctaaactggttcagtgtttg |
| **Ins. 102-103/GT** | 9 | acaagctggagtacaacggcaccatcagt ggttgcctgacagtaaaactctctg | 26 | Gctgttgtagccggcggaggttccatgttgg ctaaactggttcagtg |
| **Ins. 103-104/G** | 10 | ggcacaagctggagtacaacggtagtggtt gcctgacagtaaaactctctgc | 27 | gttgtcgctgttgtagcctgcagagatggttc catgttggctaaactggttcagt |
| **Ins. 104-105/GT** | 11 | acaagctggagtacaacggcaccggttgc ctgacagtaaaactctctgcc | 28 | Gctgttgtagccggcggaactgatggttcca tgttggctaaactggtt |
| **Ins. 103-105/G** | 12 | ggcacaagctggagtacaacggtggttgcc tgacagtaaaactctctgcc | 29 | gttgtcgctgttgtagcctgcagagatggttc catgttggctaaactggttcagt |
| **Ins. 103-105/GT** | 13 | acaagctggagtacaacggcaccggttgc ctgacagtaaaactctctgcc | 30 | Gctgttgtagccggcggagatggttccatgtt ggctaaactggttcagt |
| **Ins. 102-105/G** | 14 | ggcacaagctggagtacaacggtggttgcc tgacagtaaaactctctgcc | 31 | gttgtcgctgttgtagcctgcagaggttccat gttggctaaactggttcagtg |
| **Ins. 102-105/GT** | 15 | acaagctggagtacaacggcaccggttgc ctgacagtaaaactctctgcc | 32 | Gctgttgtagccggcggaggttccatgttgg ctaaactggttcagtg |
| **C106S** | 16 | ggcaccggtagcctgacagta | 33 | tactgtcaggctaccggtgcc |
| **NemR edges (PCS2)** | 17 | atatatatcgatgccaccatgaacaaacac accgaacatgatactcgc | 34 | Atatattctagactaaacggcaggcgtcgca ataatgtttttac |
